# Tau catalyzes amyloid-β aggregation in a fold-dependent manner

**DOI:** 10.1101/2025.10.21.683640

**Authors:** Michele Mosconi, Chiara Leonardi, Zev Armour-Garb, Beatrice Rocutto, Marten Beeg, Georg Meisl, Lei Ortigosa-Pascual, Luca Broggini, Mario Salmona, Stefano Ricagno, Tuomas P. J. Knowles, Luisa Diomede

## Abstract

Interactions between amyloidogenic proteins are emerging as critical drivers of neurodegenerative diseases, yet the molecular mechanisms remain poorly understood. In Alzheimer’s disease (AD) and chronic traumatic encephalopathy (CTE), co-deposition of tau and amyloid-β (Aβ) leads to accelerated disease progression. Here, we investigated the direct interaction between Aβ and tau, combining *in vitro* reconstruction, computational modeling, and *in vivo* models. We show that tau aggregates with AD paired helical filament (PHF) and CTE folds catalyze the primary nucleation of Aβ42 in a fold-specific manner, through an enzyme-like kinetic with recognition mechanisms. CTE fibrils exhibit the highest catalytic activity, also constraining Aβ42 polymorphism. PHF and CTE tau fibrils increase Aβ42 toxicity in SH-SY5Y neuroblastoma cells and transgenic *Caenorhabditis elegans*. Catalytic heterotypic interactions between amyloidogenic proteins offer new insights into the pathological mechanisms of multiple proteinopathies. The mechanisms described here may guide the structure-based design of new therapeutic agents targeting specific amyloidogenic interactions.

## Introduction

Amyloid formation is a pathological hallmark of several neurodegenerative diseases, including Parkinson’s Disease, Amyotrophic Lateral Sclerosis, Alzheimer’s disease (AD), and Chronic Traumatic Encephalopathy (CTE)^1–4^. Amyloid fibrils are formed by misfolded proteins that aggregate with characteristic cross-β structures via nucleation-dominated pathways, namely primary and secondary nucleation^5^. Numerous proteins, such as α-synuclein, tau, and amyloid-β (Aβ), aggregate via secondary nucleation-dominated mechanisms^6–8^, which is reported to occur on fibril’s structural defects and is responsible for the catalytic amplification of amyloids^9,10^.

For amyloidogenic proteins such as α-synuclein, tau, and TAR DNA-binding protein 43 (TDP-43), the process of amyloid assembly leads to the formation of disease-specific polymorphs, suggesting that pathological conditions control aggregation processes^11–15^. Among these, tauopathies represent a striking case of disease-specific polymorphism^11,15^. Most AD-associated tau fibrils adopt a paired helical filament (PHF) structure^16^, whereas CTE fibrils exhibit looser helical twists with greater surface area^17^. Yet the role and mechanisms underlying amyloid polymorphism in disease conditions remain largely unknown. Looking into the role of tau polymorphism will clarify how tau aggregation processes might shape disease onset and progression.

Neurodegenerative disorders commonly comprise multiple proteinopathies, characterized by aggregation of different proteins into amyloid fibrils^18,19^. Within this milieu, interactions between heterogeneous amyloidogenic proteins are increasingly recognized as potential drivers of disease progression^20,21^. Several cross-seeding mechanisms have been studied at the molecular level, including Aβ42-Aβ40 and Aβ42-⍺-synuclein^22,23^. Tau-Aβ crosstalk remains largely uncharacterized due partly to the poor mechanistic understanding of spontaneous tau aggregation in disease conditions^24,25,7^. Nonetheless, *in vivo* evidence has demonstrated that tau deletion protects against Aβ-mediated toxicity, suggesting that tau may be involved upstream of Aβ pathology^26^. Moreover, the observation of tau-Aβ co-localization at diseased synaptic terminals^27^ and association of intracellular Aβ to neuronal vulnerability^28^ suggest the presence of an extensive landscape for tau and Aβ interaction, encompassing both the intra- and extracellular environment. These interactions may be particularly relevant in AD and severe forms of CTE, where both tau and Aβ deposition occur, suggesting a key role in the progression of the pathology and neurodegeneration^27,29^. Studying the crosstalk between aggregation-prone proteins at a mechanistic level might highlight key interactions linking the co-deposition of multiple proteins with neuronal death and neurodegeneration.

Here, we report the first mechanistic description of tau-Aβ crosstalk in disease-mimicking conditions. Kinetics, biochemical, and biophysical approaches were applied to investigate the interaction of tau fibrils with PHF and CTE folds with Aβ. In particular, we used a tau fragment spanning the amyloid core residues (297-391) and capable of spontaneously assembling into amyloids with PHF and CTE folds^24,30^ (hereafter referred to as tau), and the Aβ42 peptide, an isoform associated with pathological conditions^6^. We found that both tau polymorphs promote Aβ42 heterotypic primary nucleation via direct interaction, exhibiting enzyme-like kinetics^31^. Tau acts with fold-specific reactivities, with CTE fibrils exhibiting higher catalytic efficiency than PHFs. Structure-reactivity comparison and docking analysis identified a specific pocket responsible for Aβ42 heterotypic primary nucleation. The pocket is highly exposed in CTE fibrils and hindered in PHF, explaining the fold-specific behaviour. The enzyme-like behaviour of such structures contrasts with nonspecific secondary nucleation, which occurs on fibril defects^9,10^.

We also investigated the biological relevance of this cross-reactivity using, as experimental models, human neuroblastoma SH-SY5Y cells treated with Aβ42 peptide alone or together with tau fibrils, and by administering tau fibrils to transgenic *C. elegans* CL2120 strain constitutively expressing Aβ42 in the body-wall muscle^32^. Tau fibrils with PHF and CTE folding increased Aβ42 toxicity both *in vitro* and *in vivo* in a fold-dependent manner.

The identification of specific mechanisms responsible for heterotypic catalytic amplification of amyloids opens new insights into the pathological processes underlying tauopathies and neurodegenerative disease. It provides a new framework for the rational development of multi-target treatments targeting specific amyloid-amyloid interactions.

## RESULTS

### Tau fibrils catalyze Aβ42 aggregation in a fold-dependent manner

Initially, we performed bulk kinetic assays with logistic fitting to determine the role of tau amyloids on Aβ42 aggregation. First, to benchmark the assay, we validated the self-aggregation of Aβ42 using Thioflavin T (ThT) as amyloid reporter dye. Kinetic modelling allowed to dissect the microscopic molecular events underlying the overall aggregation process^33–35^. We determined secondary nucleation as the dominant microscopic mechanism guiding the overall aggregation process, consistently with previous reports^6^ **(Supplementary Fig.1A**). Nucleation-elongation kinetic models, not accounting for secondary pathways, and fragmentation-dominated models resulted in misfits^6^ (**Supplementary Fig.1B, C**). We further validated this analysis by performing power exponent (𝛾) analysis of the half-time (t_1/2_) as a function of initial monomer concentration (m_0_) following the relationship 𝑡_!/#_∼𝑚^%36^. We determined a scaling exponent 𝛾 = −1.39 ± 0.05, consistent with secondary nucleation-dominated aggregation as reported by *S. I. A. Cohen et al.*^6^ (**Supplementary Fig. 1D**). Kinetic rate constants for primary nucleation, secondary nucleation, and fibril elongation of Aβ42 were determined by global fitting (**Supplementary Fig. 1E**). Within the range of 3 - 6 µM the aggregation is strongly dependent on monomer concentration; therefore no saturation of secondary nucleation was observed, suggesting that the threshold concentration is above 6 µM^6^.

Disease-specific tau fibrils designed to mimic PHF and CTE folds were produced as reported in the literature employing the tau fragment spanning residues 297-391^24,30^, and fibrillar morphology was confirmed by transmission electron microscopy (TEM) (**Supplementary Fig. 2A-G**).

The effect of PHF and CTE seeds on Aβ42 aggregation (**Fig. 1A, B**) was determined by monitoring ThT fluorescence for 15 hours. The data were normalized to the seed’s baseline fluorescence to isolate the effect of Aβ42 changes in fluorescence due to aggregation. Tau seeds alone produced a constant ThT signal over this timescale, indicating that fluorescence increases should originate from Aβ42 aggregation (**Supplementary Fig. 2H**). By fitting the data with the logistic function (Eq. 1)^37^ described in Methods, we observed that both PHF and CTE seeds conferred a concentration-dependent decrease in lag time (t_lag_)^38^ of Aβ42 aggregation (**Supplementary Fig. 3A**). Remarkably, fold-specific differences between the two tau polymorphs also emerged. CTE seeds significantly reduced the t_lag_ at 0.125 µM, corresponding to a seed concentration as low as 3% of the total Aβ42 protein, whereas the effect of PHF became significant from 0.5 µM. These effects at sub-stoichiometric tau:Aβ42 molar ratios suggest that tau fibrils can exert a catalytic effect on Aβ42 aggregation, similarly to catalytic secondary nucleation events in self-aggregation reactions^33–35^.

**Figure 1.**
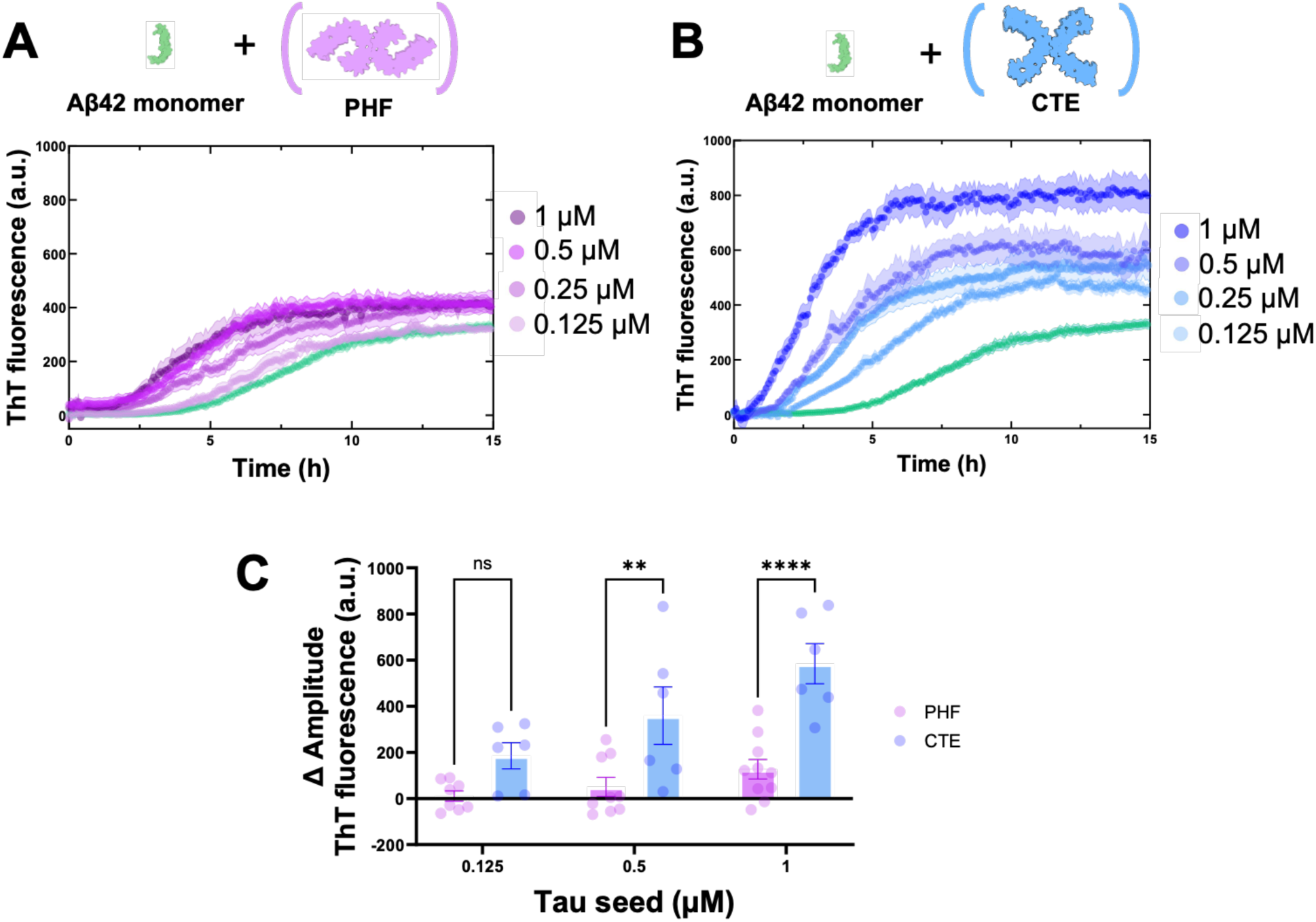
Concentration-dependent effect of PHF and CTE tau seeds on Aβ42 aggregation. **A, B**. ThT fluorescence profiles of 4 µM monomeric Aβ42 aggregation in the absence (green line) or presence of different concentrations of **A.** PHF (purple lines) and **B.** CTE (blue lines) tau seeds. Curves derived from the analyses of the ThT fluorescence, expressed as fluorescence arbitrary units (a.u.) obtained at different time-points and expressed as the mean ± SEM (N=3) from at least three independent experiments. **C.** Variation of ThT fluorescence amplitude (ΔA) determined from the data represented in panels **A** and **B,** calculated as ΔA=A_Aβ42+tau_-A_Aβ42_. Data are the mean ± SEM (N=3) from at least three independent experiments. **** p<0.0001, ** *p*<0.01, and ns = not significant according to two-way ANOVA and Tukey’s *post hoc* test. Both PHF and CTE seeds accelerate Aβ42 aggregation, increasing ThT amplitude in a fold-specific manner.

PHF and CTE seeds also elicited concentration-dependent increases in ThT amplitude. CTE exhibited the strongest of such effects, with an approximately two-fold increase at the highest tested concentration (**Fig. 1C**). These data suggest that tau fibrils promote Aβ42 aggregation in a fold-dependent manner, with polymorph-specific structural features conferring distinct catalytic activities.

### Tau amyloids catalyze heterotypic Aβ42 primary nucleation

In order to identify the microscopic mechanisms driving the catalytic effect of tau seeds on Aβ42, we monitored the aggregation of different Aβ42 concentrations in the presence of PHF or CTE seeds (**Fig. 2A, B**). The 𝛾 value of −1.39 ± 0.05 determined for Aβ42 self-aggregation (**Supplementary Fig. 1D**) rose to −0.6 ± 0.2 and −0.4 ± 0.2 in the presence of PHF and CTE fibrils, respectively (**Fig. 2C**). These values approach the theoretical limit of −0.5, indicating that the dominant microscopic mechanism is saturated and independent of the Aβ42 monomer concentration^39^. CTE seeds decrease the t_1/2_ of Aβ42 aggregation more than PHFs (**Fig. 2C**), further confirming the fold-specific catalytic activity of tau polymorphs (**Supplementary Fig. 3B**).

**Figure 2.**
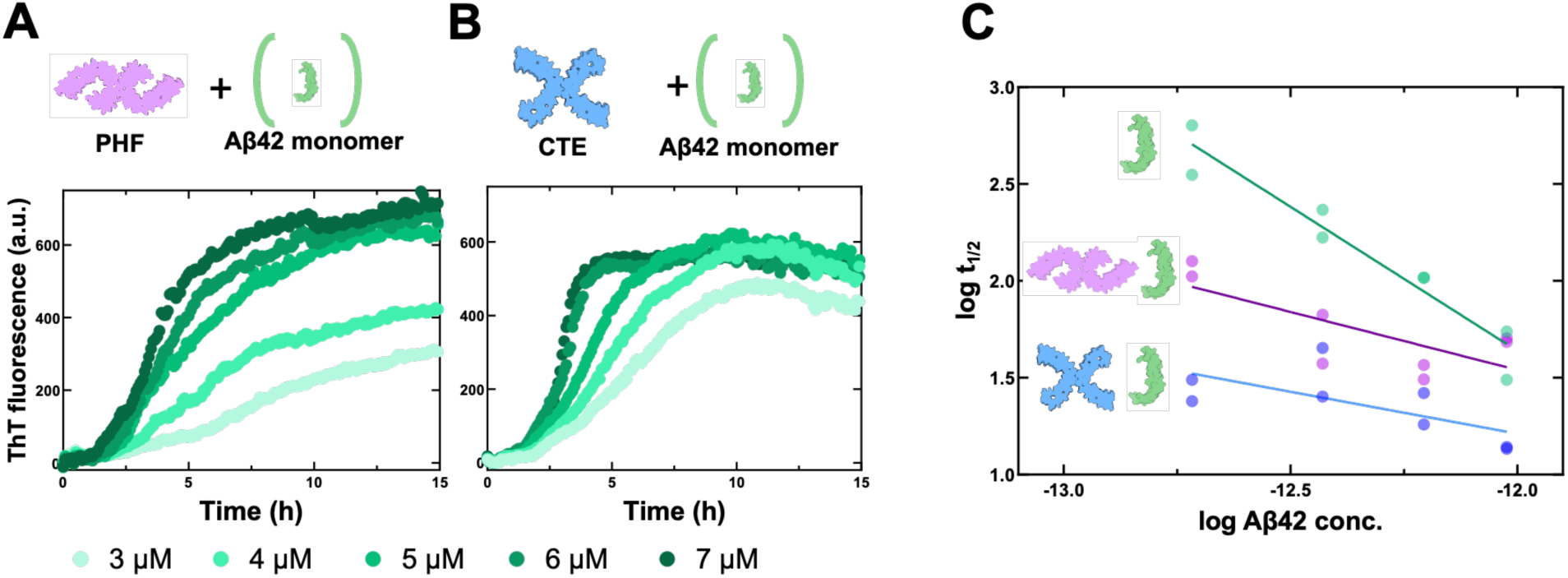
Heterotypic Aβ42 aggregation with PHF- and CTE-seeds is dominated by saturated mechanisms. **A-B.** ThT fluorescence profiles of 0.25 µM **A.** PHF- and **B.** CTE-seeded aggregations of different Aβ42 concentrations (3 - 7 µM). Curves derived from the analyses of the ThT fluorescence, expressed as arbitrary units (a.u.), were obtained at different time-points and expressed as the mean of at least three independent experiments. **C.** Double logarithmic plot of half-time (t_1/2_) *vs.* Aβ42 concentration, allows extrapolation of the power exponent (𝛾) for PHF-seeded (purple) and CTE-seeded (blue) Aβ42 aggregations compared to the power exponent of Aβ42 self-aggregation (green). Data from two independent experiments are plotted for each condition.

We performed kinetic modeling to characterize the microscopic kinetics of tau-mediated Aβ42 aggregation. Kinetic curves obtained at different Aβ42 monomer concentrations in the presence of fixed amount of PHF and CTE seeds (**Fig. 2A, B**) were normalized and globally fitted as described in Methods^33–35^. The data for both PHF- and CTE-seeded reactions were best fitted by a saturated secondary nucleation model (**Fig. 3A**), wherein saturation arises from reversible monomer-fibril binding followed by irreversible formation of elongation-prone nuclei^39^. Failing to account for saturation effects resulted in a misfit (**Fig. 3B**). PHF- and CTE-seeded aggregations exhibited a saturated mechanism at monomer concentrations as low as 3 µM, whereas in self-aggregation conditions saturation occurs at concentrations greater than 6 µM as mentioned before (**Supplementary Tab. 1A**).

**Figure 3.**
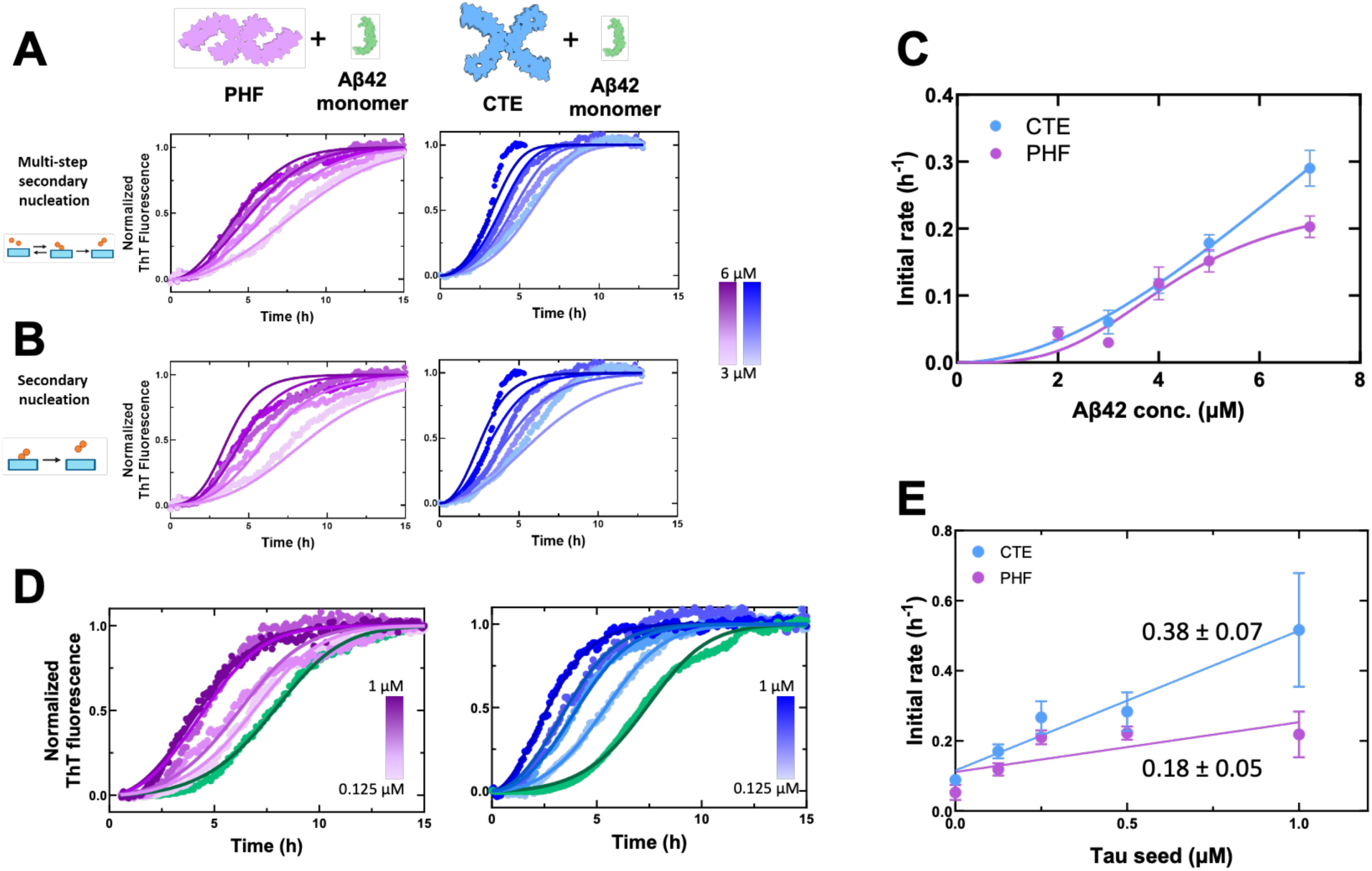
Tau amyloids catalyze Aβ42 aggregation with enzyme-like kinetics. **A-B.** Global fitting analysis to microscopic kinetic equations obtained from the normalization of ThT fluorescence values shown in Figure 2A and B, in which the aggregation of 3-6 µM Aβ42 in the presence of 0.25 µM PHF- or CTE-seeds was evaluated. **A.** Global fitting of Aβ42 aggregation in the presence of PHF and CTE tau seeds with multi-step secondary nucleation resulted in the best fit. In contrast, **B.** secondary nucleation models resulted in a misfit. Data are the mean (N=3) of 3 independent experiments. **C.** The initial rate of fibril formation was calculated for the aggregation of different Aβ42 concentrations (2-7 µM) seeded with 0.1 µM tau seeds, the data were fitted to an enzymatic kinetic model accounting for cooperative binding of Aβ42 to tau fibrils, as described in Methods. Positive cooperativity with n > 1 was determined in both PHF- and CTE-seeded conditions. Data are mean ± SEM (N = 3). **D.** Global fitting analysis to microscopic kinetic equations from the normalization of ThT fluorescence values shown in Figure 1A and B of 4 µM Aβ42 in the presence of increasing PHF or CTE seed concentrations (0.125 - 1 µM). Aβ42 self-aggregation is shown in green. Data are the mean (N=3) of three independent experiments. The microscopic parameters determined by global fitting in **A.** fitted optimally the kinetic curves, further confirming the microscopic kinetic analysis. **E.** The initial rate values were calculated for each seed concentration tested (0.125-1 µM) on 4 µM Aβ42, by global fitting with a multi-step secondary nucleation model. The catalytic activity of PHF and CTE fibrils was assessed by fitting a regression line to the data. Data are mean ± SEM (N=3). Slope of lines are mean ± SEM.

To further explore the catalytic activity of tau, we determined the initial rate of fibril formation^6^ (𝜆 ∝ 𝑘_n_𝑘_+_) for different tau:Aβ42 molar ratios from 1:30 to 1:70. At ratios higher than 1:70, Aβ42 self-aggregation dominated, obscuring any effect of tau. We observed a sigmoidal trend in PHF-seeded conditions in the range of concentration tested, whereas in the presence of CTE seeds at 1:70 ratios, the plateau was still not reached. These data suggested that CTE seeds greatly affected initial rates of Aβ42 aggregation, also at high tau: Aβ42 ratios, whereas PHFs do not, highlighting the greater catalytic activity of CTE polymorphs (**Fig. 3C**). Remarkably, fitting to an enzyme-like model (Hill equation)^31^ (Eq. 2) captured the kinetics and uncovered positive cooperativity for both PHF and CTE, consistent with Aβ42-mediated nucleation on tau fibrils (**Fig. 3C**).

To quantify the efficiency of tau seeds in catalyzing Aβ42 primary nucleation, we calculated the value of λ at varying seed concentrations. The k_+_ was held constant for all the conditions, while k_n_ was fitted for each seed concentration (**Fig. 3D** and **Supplementary Tab. 1B**). In the absence of seed, the λ value of ∼ 0.1 h^−1^ was consistent with our Aβ42 self-aggregation kinetic model. Linear regression of λ against tau seed concentration produced slopes of 0.18 ± 0.05 (PHF) and 0.38 ± 0.07 (CTE), with the larger CTE slope indicating stronger catalysis of Aβ42 heterotypic primary nucleation (**Fig. 3E**). Hence, the differential structural properties of PHF and CTE folds result in altered catalytic effects on Aβ42 aggregation.

To understand the origins of such a catalytic mechanism, we combined comparative structural analysis and *in silico* docking experiments. PHF and CTE fibrils differently expose residues 351-378, constituting an optimal nucleation site for Aβ42 (**Supplementary Fig. 4A**). Flexible docking analysis revealed the ability of the subregion 353-369 to bind different portions of Aβ42 monomeric units (**Supplementary Fig. 4B-D**). Core regions of Aβ42 yielded greater scores compared to N-ter and C-ter regions (**Fig. 4A, B**) in accordance with previously proposed interacting surfaces^40,41^. This surface is exposed to the solvent in the CTE fold, whereas in PHF, the high steric hindrance reduces its accessibility, plausibly explaining the fold-specific reactivity. The atomistic structural determinants of such transient catalytic interactions between amyloidogenic proteins are yet to be clarified.

**Figure 4.**
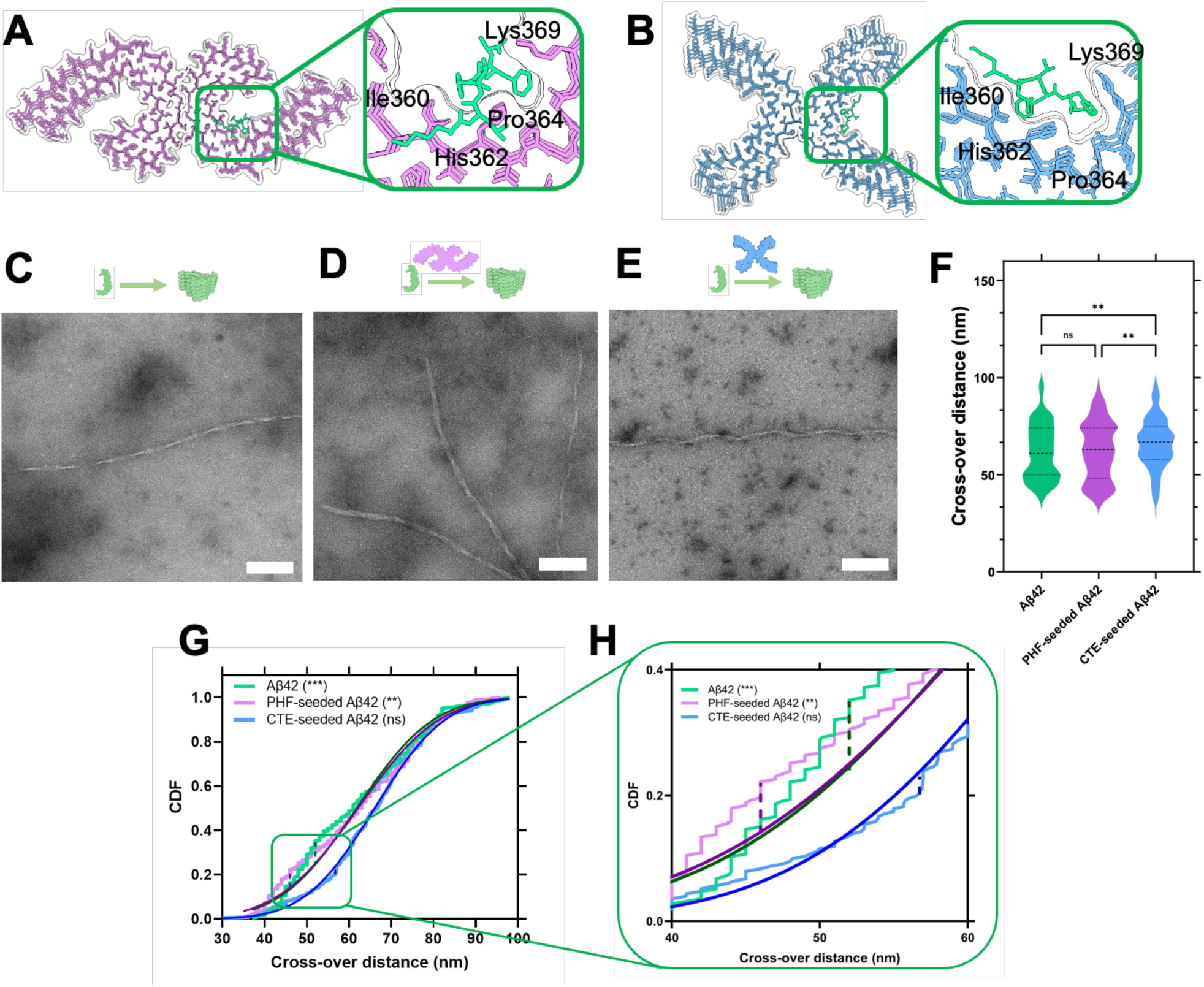
Characterization of Aβ42 amyloid fibrils formed in the presence and absence of tau seeds. Molecular docking of the core region of Aβ42 (KLVFFA) (green) on **A.** PHF (PDB: 8Q8R) and **B.** CTE (PDB: 6NWP) fibrils. A specific catalytic surface spanning residues 360-369 of tau amyloids was identified. **C-E**. Representative images obtained from transmission electron microscopy analysis performed by negative staining of fibrils produced 15 h after **C.** self-aggregation of 4 µM Aβ42 alone and in the presence of **D.** 0.25 µM PHF or **E.** CTE seeds. Scale bar = 100 nm. **F.** Cross-over distances of Aβ42 fibrils formed in self- and cross-seeded reactions. Data are shown as a violin plot of cross-over distance measurements from two independent experiments (N > 100). Dotted lines are the median, first, and third quartiles. \*\**p*<0.01, ns = not significant according to two-way ANOVA and Tukey’s *post hoc* test. **G.** Empirical cumulative distribution functions (CDFs) of crossover distances for Aβ42 fibrils (green), PHF-seeded (purple), and CTE-seeded (blue) Aβ42 fibrils. **H.** Enlarged view of dotted lines representing the reference cumulative distribution shown in panel **G**. \*\**p*< 0.01 and \*\*\**p*< 0.001 *vs*. normal distribution for PHF-seeded Aβ42 and Aβ42, respectively, according to one-sample Kolmogorov–Smirnov test. CTE-seeded Aβ42 fibrils do not significantly differ from a normal distribution.

### Tau catalytic activity affects Aβ42 polymorphism

Increasing concentrations of tau seeds led to an increase in ThT fluorescence signal of Aβ42 up to 50% (**Fig. 1C**), suggesting that tau fibrils promote the formation of either more Aβ42 fibrils or Aβ42 polymorphs with higher affinity to ThT. Nevertheless, such increase in ThT fluorescence cannot be accounted for solely by increased fibril mass concentration, given the solubility of Aβ42^37^. To determine whether this observation reflects structural changes, we examined the morphology of Aβ42 fibrils formed by self-aggregation in PHF- and CTE-seeded conditions by TEM analysis (**Fig. 4C-E**). Analysis of fibril cross-over distances revealed statistically significant differences among unseeded, PHF- and CTE-seeded conditions (**Fig. 4F**). In unseeded and PHF-seeded conditions, fibrils displayed a similar broad distribution of cross-over distances from 30 to 100 nm, consistent with structural polymorphism. Conversely, CTE-seeded Aβ42 fibrils exhibited a narrower distribution centred around 66 nm, indicative of more uniform fibril morphology (**Supplementary Fig. 5A**). Statistical analysis revealed that the distribution of cross-over distances in the presence of CTE seeds differed from the other two aggregation conditions (**Supplementary Fig. 5B**). CTE-seeded crossover distances did not differ significantly from a normal distribution; by contrast, PHF-seeded and self-aggregated Aβ42 fibrils showed significant deviations, with the latter exhibiting the largest maximum deviation (**Fig.4G**). Taken together, these data suggest a constrained polymorphism of Aβ42 fibrils in the presence of CTE seeds, with PHF seeds originating an intermediate scenario. These measurements were not determined to have been significantly confounded by tau seeds, as indicated by pairwise analyses and two-sample Kolmogorov-Smirnov (KS) test of the Aβ42 fibril width distribution during self-aggregation compared to tau-seeded aggregations (**Supplementary Fig. 5C-E**). Furthermore, we observed a reduction of fibril length under cross-seeding conditions, suggesting that tau fibrils accelerated the rate of nucleation and decreased the free Aβ42 monomer concentration available for fibril elongation (**Supplementary Fig. 5F**).

Taken together, morphological analyses suggest that tau fibrils with a CTE fold constrain Aβ42 polymorphism. We speculate that the PHF seed preparation exerts a similar but weaker effect, not readily detectable by TEM analysis, consistent with its lower impact on ThT fluorescence amplitude. These findings support the presence of reactive catalytic-like surfaces on tau amyloids that not only accelerate Aβ42 nucleation but also template specific amyloid assemblies.

We assessed the presence of interaction between Aβ42 and tau amyloids by immunoprecipitation with an antibody against human Aβ (**Supplementary Fig. 6**). In cross-aggregation conditions, tau co-precipitated with Aβ42, highlighting a strong direct interaction in accordance with their *in vivo* co-localization at the synaptic level^27^.

### Disease-relevant tau polymorphs catalyze Aβ42 nucleation selectively

To assess whether disease-relevant tau fibrils selectively catalyze Aβ42 nucleation, we prepared disease-unrelated tau assemblies using heparin, the polyanion most used and reported to induce the formation of amyloid not associated with tauopathies^42^ (**Fig. 5A**). Although a reduction of t_1/2_ of Aβ42 was observed upon addition of heparin-tau seeds, the effect was variable across replicates and not concentration-dependent (**Fig. 5B**). This indicates that the negatively charged surfaces contained in tau seeds and increased surface area alone cannot account for the catalytic activity observed for PHF and CTE fibrils. Intrinsic structural and chemical properties of disease-relevant tau amyloids are therefore responsible for the selective catalytic activity toward Aβ42 primary nucleation. To further validate the selectivity of the catalytic process, we used fibrils of Atrial Natriuretic Peptide (ANP), characteristic of isolated atrial amyloidosis (IAA)^43^, which is unrelated to tauopathies. ANP amyloids prepared as described in Methods were further used to seed Aβ42 aggregations (**Fig. 5C**). No statistically significant change in t_1/2_ at each seed concentration tested was observed, suggesting that ANP amyloids do not act as a catalyst for Aβ42 heterotypic primary nucleation (**Fig. 5D**).

**Figure 5.**
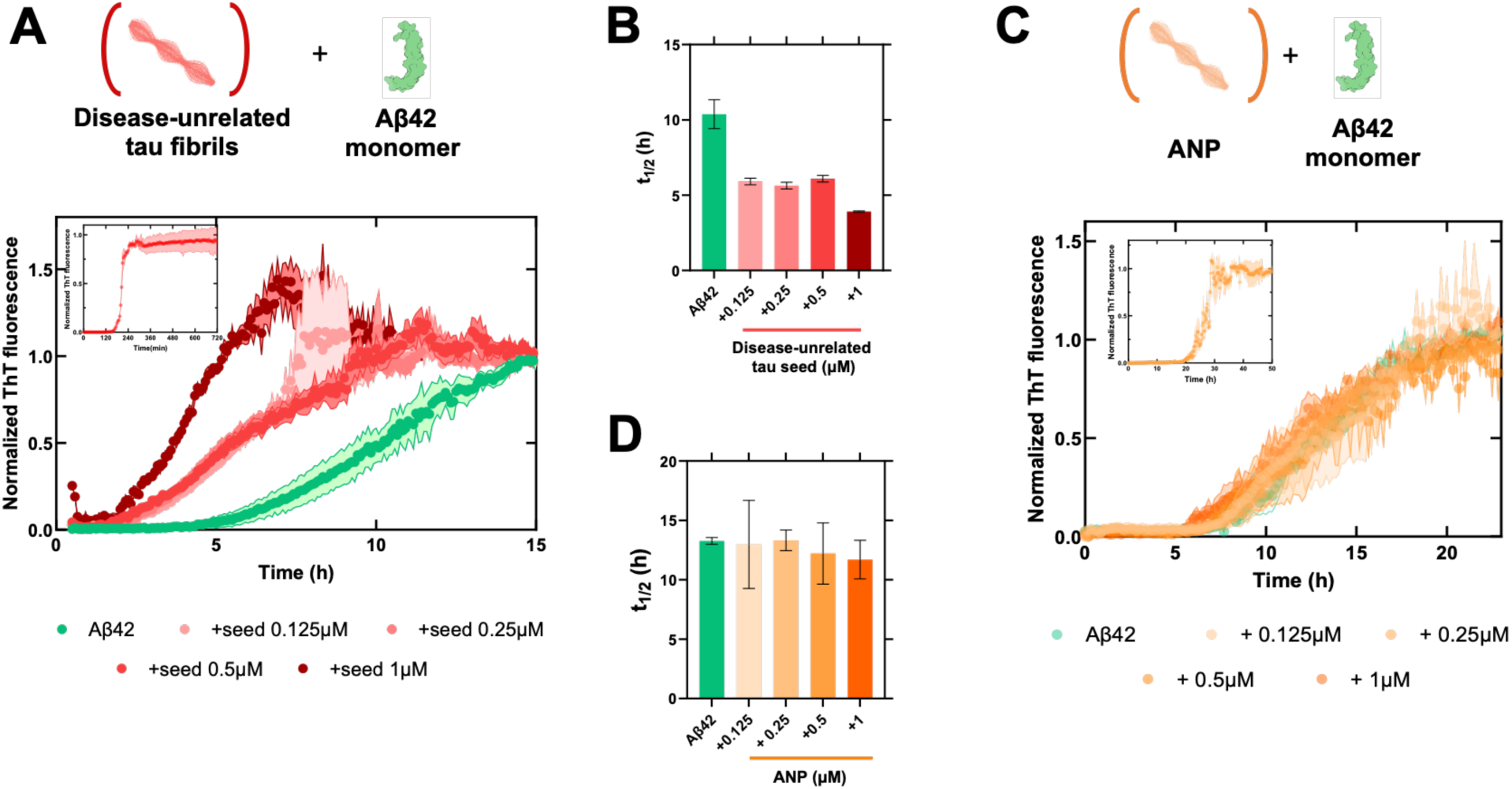
Specific determinants control tau-Aβ42 crosstalk. **A.** Effect of disease-unrelated tau amyloids produced in the presence of heparin on Aβ42 aggregation. Normalized ThT fluorescence of 4 µM monomeric Aβ42 aggregation in the absence (green line) or presence of different concentrations of disease-unrelated tau seed. Insert is a representative tau amyloid assembly reaction. The first 15 hours of the aggregation are shown. **B.** Analysis of t_1/2_, extrapolated by logistic fitting of Aβ42 aggregation in the absence or presence of increasing concentration of tau amyloids produced in the presence of heparin. **C.** Effect of ANP amyloid on Aβ42 aggregation. Normalized ThT fluorescence of 4 µM monomeric Aβ42 aggregation in the absence (green line) or presence of different concentrations of ANP seeds (yellow). The insert is a representative ANP assembly reaction. The first 24 hours of the aggregation are shown. ANP amyloids do not affect the aggregation of Aβ42; the kinetic curves are superimposable, and no shift in lag or half-time is detected. **D.** Analysis of t_1/2_, extrapolated by logistic fitting of Aβ42 aggregation in the absence or presence of increasing concentration of ANP. Data are mean ± SEM of at least three independent experiments.

These findings suggest that PHF and CTE polymorphs act as selective catalysts for Aβ42 nucleation, and that structural and physicochemical determinants are required for an effective tau-Aβ42 crosstalk.

### Tau polymorphs affect the biological activity of Aβ42

We next asked whether PHF- and CTE-mediated cross-seeding of Aβ42 could impact its biological *in vitro* and *in vivo* activity. In SHSY-5Y cells, 24 hours of exposure to 1.25-5 µM PHF seeds alone did not affect the viability, whereas CTE seeds reduced viability by ∼45% at all tested concentrations (**Supplementary Figure 7**). The greater cytotoxicity of CTE versus PHF may reflect the higher hydrophobicity of CTE seeds. Cells were then treated for 24 hours with monomeric Aβ42 alone or together with PHF and CTE seeds, maintaining the Aβ42:tau molar ratio used in aggregation experiments (**Fig. 6A**). Under these conditions, Aβ42 reduced cell viability by 19% (100% ± 1.8 and 80.78% ± 3.8 viability for vehicle- and Aβ42-treated cells, *p*<0.0001), in accordance with our previous observation^44^. The co-administration of Aβ42 with PHF and CTE tau seeds worsened its toxicity by 15% and 54%, respectively (53.20% ± 3.0 and 37.23% ± 1.4 viability for Aβ42 + PHF and CTE seeds, respectively, *p*<0.0001) (**Fig. 6B-C**). The co-occurrence of Aβ42 and tau polymorphs resulted in a ∼24% stronger reduction in viability than expected from independent additive effects, consistent with synergistic toxicity.

**Figure 6.**
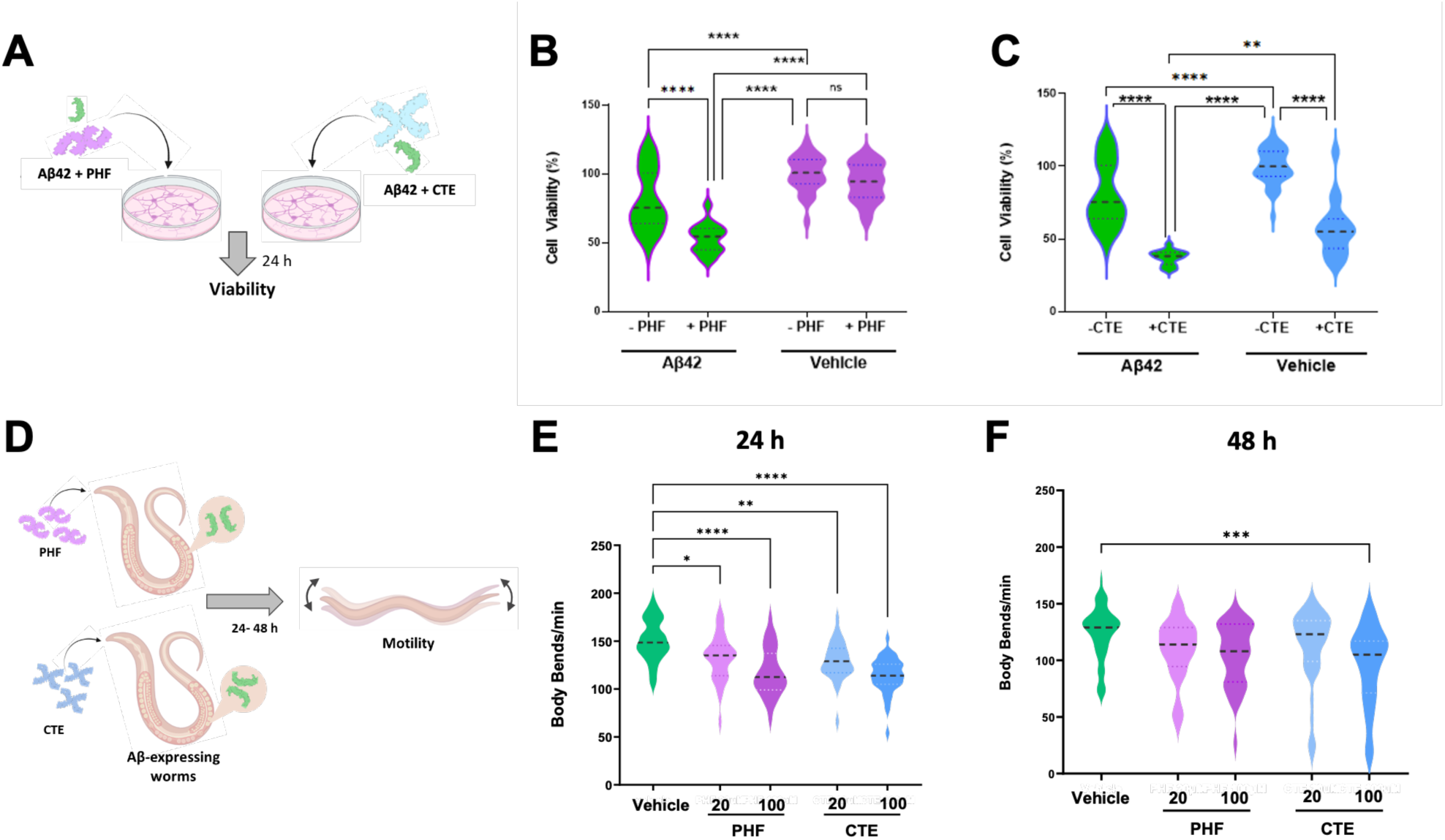
Tau seeds impact the biological effect of Aβ42 *in vitro* and *in vivo*. **A.** Representative scheme of the *in vitro* experiments. SHSY-5Y neuroblastoma cells were treated with 10 µM monomeric Aβ42 alone or with **B.** 1.25 µM PHF or **C**. CTE seeds, and the cell viability was determined after 24 hours. Control cells were treated with the medium alone (Vehicle). Data are expressed as the mean ± SD of % viability compared to Vehicle-treated cells. N = ∼40 from 7 independent experiments. ** *p*<0.05 and \*\*\*\**p*<0.0001 according to two-way ANOVA and Bonferroni *post hoc* test. Interaction Aβ/PHF *p* = 0.006 and Aβ/CTE interaction *p* = 0.97. **D**. Representative scheme of the *in vivo* experiments. Synchronized transgenic CL2120 worms expressing Aβ42, at the L3 larval stage, were fed 20 and 100 µM PHF and CTE seeds suspended in 50 µL of aggregation buffer. Control worms were fed 5 mM PBS, pH 7.4 (Vehicle). **E.** Twenty-four hours and **F**. 48 hours after the administration, the motility of worms was determined by scoring the number of body/bends per minute (body/bends/min). Data are the mean ± SEM (N = 30) from 3 independent experiments. * *p*<0.5, ** *p*<0.05, *** *p*<0.0005, and \*\*\*\**p*<0.0001 according to one-way ANOVA and Bonferroni *post hoc* test. Dotted lines are the median, first, and third quartiles.

*In vivo* experiments were performed using the CL2120 transgenic *C. elegans* strain, which constitutively accumulates Aβ42 fibrils in muscle cells^32^ (**Fig. 6D**). CL2122 worms were employed as controls. Amyloid accumulation in CL2120 results in the onset of a progressive defect in their locomotion ability, scored as the number of movements in a liquid in a minute (body bends/min) (**Supplementary Figure 8A - B**)^45^. Based on our previous studies on tau-induced toxicity in nematodes^46^ the impact of PHF and CTE seeds was evaluated by administering them at 10-100 µM. Twenty-four hours after administration, PHF and CTE seeds significantly decreased CL2120 worm movement in a concentration-dependent manner (**Fig. 6E**). In particular, the body bends/min of CL2120 nematodes was significantly reduced by 12% at 20 µM and by 22% at 100 µM in PHF-treated worms, and by 14% and 24% in 20 µM and 100 µM CTE-treated worms, respectively (**Fig. 6E**). After 48 hours CTE, but not PHF, administered at the highest concentration, reduced the worms’ motility of 24% (**Fig. 6F**). At these time points, the motility of CL2122 remained unaffected by treatment with PHF and CTE seeds (**Supplementary Fig. 8C, D**). These data ruled out simple additive toxicity between Aβ42 and tau, demonstrating instead that tau enhances Aβ42 toxicity in a fold-specific manner, with CTE fibrils exerting a greater effect. Different tau polymorphs thus emerged as key modulators of Aβ42 toxicity both *in vitro* and *in vivo*.

## Discussion

Deposition of protein aggregates is a hallmark of many neurodegenerative diseases^1–4^. Several pathological conditions involve the accumulation of more than one amyloidogenic protein, such as Aβ, tau, α-synuclein, and TDP-43, highlighting their possible interaction in disease onset or progression^18–21^. However, the molecular mechanisms underlying the heterotypic interactions between different aggregation-prone proteins are still poorly understood.

AD is the most common form of multiple proteinopathy, accounting for 60-70% of all cases of dementia. More than 55 million people worldwide are affected by dementia, and numbers are expected to increase to 145 million by 2035^47^. Pharmacological intervention is still limited in part due to an incomplete understanding of the molecular mechanisms underlying such multiple proteinopathies and the intertwined interactions between amyloidogenic proteins. This study investigated the interaction between tau and Aβ42, two key amyloidogenic proteins involved in AD and other tauopathies such as CTE. To this end, the self-aggregation process of Aβ42 was perturbed using disease-relevant PHF and CTE tau amyloids, demonstrating that tau polymorphs behave as fold-specific catalysts of Aβ42 heterotypic primary nucleation. This catalytic interaction occurs on specific structural pockets, accelerating Aβ42 aggregation and modifying its overall aggregation process. CTE fibrils act with higher catalytic activity, constraining the structural polymorphism of Aβ42 fibrils.

Here, the observation of fold-specific reactivities for distinct tau polymorphs suggests that the region spanning the residues 351-379 of tau might act as a catalytic site for Aβ42 nucleation. Further structural analysis will allow us to characterize the catalytic site on tau amyloids responsible for the interaction with Aβ42. Previous studies have proposed the sequence KLVFFA as a possible interaction site^40,41^, and a recent study demonstrated by HMQC NMR that tau-Aβ42 interaction occurs on this region^48^. Our results are in accordance with these observations. Docking simulations suggest preferential binding of Aβ42 core regions compared to its N- and C-terminal domains, plausibly triggering heterotypic primary nucleation. In addition, direct interaction between tau seeds and Aβ42, strong enough to resist immunoprecipitation, can explain the co-localization of the two proteins observed in disease conditions^27^.

We also demonstrated that specific structural and physicochemical determinants control the fold-specific catalytic interaction observed, as demonstrated by the absence of a catalytic effect on Aβ42 nucleation of tau fibrils produced in the presence of heparin, and of amyloid fibrils produced by ANP, a protein totally unrelated to neurodegenerative disorders.

Our model of amyloid-amyloid heterotypic interaction highlights the importance of structural compatibility in heterotypic amyloid interactions. Additionally, the reduced polymorphism observed in CTE-seeded conditions aligns with a templated nucleation mechanism driven by amyloid surfaces. Whether similar mechanisms occur in other heterotypic aggregation systems remains to be determined.

These findings reveal a selective enzyme-like heterotypic nucleation mechanism underlying tau-Aβ42 interaction in disease-mimicking conditions, contrasting with classical Aβ42 secondary nucleation, which was demonstrated to occur nonspecifically at fibril surface defects^9,10^. To our knowledge, this is the first observation of a catalytic mechanism mediating cross-reactivity between distinct amyloidogenic proteins, occurring on specific surfaces.

We next showed that the heterotypic aggregation of Aβ42 in the presence of tau polymorphs influences its biological activity both *in vitro* and *in vivo*, emphasizing the pathological significance of tau-mediated Aβ toxicity. This is in line with the co-occurrence of tau and Aβ at diseased synaptic terminals^27^ and preclinical evidence of interconnected Aβ-induced neurotoxicity and tau pathology^27,28^. Notably, the higher catalytic activity and greater toxicity of CTE fibrils may reflect differences in hydrophobicity and surface polarity, consistent with the clinical observation of more severe CTE pathology in the presence of Aβ deposits^29^.

Taken together, these observations support a broader model in which neurodegeneration may proceed through interconnected double and multiple proteinopathies, evolving via heterotypic amyloid-amyloid interactions. For example, CTE pathology is also linked to Lewy body dementia, suggesting that the interaction between tau and α-synuclein may also play a role in disease progression^49^. A similar scenario was also found in amyotrophic lateral sclerosis/Parkinsonism-dementia^50^.

In conclusion, we propose that specific surfaces on tau amyloids can mediate the selective and fold-specific nucleation of Aβ42, acting as molecular templates that control both the rate and structure of resulting aggregates. The mechanism here proposed provides a molecular explanation for the spatially localized amyloid cross-interactions observed in neurodegenerative diseases. Moreover, identification of key catalytic surfaces involved in tau-Aβ heterotypic interactions, as well as uncovering polymorph-specific reactivities of tau amyloids with other amyloidogenic proteins, will offer new frameworks for the structure-based design of therapeutics acting simultaneously on multiple protein aggregation and proteotoxic events.

## Acknowledgments

We thank Prof. M. Goedert for providing the tau plasmids and Dr. H. Greer for helping with TEM micrograph acquisition. *C. elegans* and OP50 *E. coli* were provided by the GCG, which is funded by NIH Office Research Infrastructure Programs (P40 OD010440). This work was partially supported by the Fondazione Sacchetti (grant 2023-2025) to L.D., the Mario Negri Alumni Association (2025) to M.M., and by FONDAZIONE CARIPLO [grant number 2024-NAZ-0018], FONDAZIONE CARIPLO/Telethon [Telethon GJC23044], Fondazione AIRC [IG 2024 ID 30307] and University of Milano, Seed 4 Innovation 2024 grant to Nano-Detox to S.R.

## Author contributions

Conceptualization, L.D., M.M., and M.B.; methodology, L.D., M.M., M.B., B.R., C.L., G.M., L.B. and Z.A.G.; formal analysis. L.D., M.M., G.M., L.O.P. and Z.A.G.; resources, L.D.; data curation, L.D., M.M., and Z.A.G.; writing—original draft preparation, L.D., M.M., C.L., L.O.P, Z. A. G., S.R., M.S.; supervision, L.D., T.P.J.K., and S.R.; funding acquisition, L.D., and S.R. All authors have read and agreed to the published version of the manuscript.

## Methods

### Aβ42 self-aggregations and kinetic modeling

Recombinant Aβ42 treated with HFIP (rPeptide, Watkinsville, GA, USA) was resuspended in 50 mM NaOH at 1 mg/mL and stored at −80°C. The compounds for which the origin is not specified were purchased from Merck (Darmstadt, Germany). Aggregation was carried out in 20 mM phosphate buffer (PB) solution, pH 8, containing 200 µM EDTA, 0.02% NaN_3_, 5 mM NaOH, and 2 µM Thioflavin-T (ThT, Merck, Darmstadt, Germany) in quiescent conditions at 37°C^6^. Six different Aβ42 concentrations ranging from 2 µM to 6 µM were screened. Seeded aggregation of Aβ42 was performed by measuring the aggregation profiles of three different monomeric protein concentrations between 3-5 µM supplemented with Aβ42 seeds at concentrations ranging from 30 to 50% of the highest protein concentration in the dilution series. Seeds were prepared by aggregation of 20 µM Aβ42 in 20 mM PB solution, pH 8, containing 200 µM EDTA, 0.02% NaN_3_, 5 mM NaOH, and 2 µM ThT at 37°C in quiescent condition^6,33^. After 5 h, the plateau equilibrium phase was reached, and aggregation mixtures were bath sonicated at room temperature for 5 min and further used to seed the Aβ42 aggregation. Aggregation assays were performed at 37°C in 96-well half-area low bind plates (3008, Corning, Corning (NY), USA), placed in a Tecan Infinite F200 (Tecan, Männedorf, Switzerland) equipped with 448 nm excitation and 485 nm emission filters. ThT fluorescence was monitored for 15 hours. At least three independent experiments and three replicates for each experiment were performed. Kinetic modelling was performed by globally fitting the datasets to a model accounting for primary nucleation, elongation, and secondary nucleation^2,3^ with the AmyloFit Platform (https://amylofit.com)^33^.

### Tau297-391 expression and purification

For expression of tau297-391 fragment^24,30^, the protein encoding vector pET3a-tau297-391 (provided by Prof. M. Goedert, MRC LMB, Cambridge) was transformed into chemically competent expression strain *E. coli* BL21(DE3). Cells were cultured at 37°C in 2xTY supplemented with 5 mM MgSO_4_ and 100 µg/mL Ampicillin. Expression was induced with 1 mM IPTG at O.D_600_ ≈ 0.8. Four hours after the induction, bacteria were harvested by centrifugation at 1400 x g for 20 min at 4°C. The pellet was suspended at 0.1 g/mL with a 50 mM MES lysis buffer, pH 6.0, containing 10 mM EDTA, 10 mM dithiothreitol (DTT), 0.1 mM 4-(2-aminoethyl) benzenesulfonyl fluoride hydrochloride, and cOmplete protease inhibitor cocktail. Cells were lysed with probe sonication (Bandelin sonopuls HD 2070, Berlin, Germany) for 7 min (5 sec on/ 10 sec off) at 95% amplitude. Lysed cells were centrifuged at 20.000 x *g* for 40 min, at 4°C. The supernatant was filtered through 0.22 μm filters and loaded into an FPLC AktaPure25M (Cytiva, Marlborough, Massachusetts, USA) equipped with HiTrap Capto S column (Cytiva, Marlborough, Massachusetts, USA) equilibrated with washing buffer (50 mM MES at pH 6.0, 10 mM EDTA, 10 mM DTT). Protein was then eluted with a linear gradient of washing buffer supplemented with 0 up to 500 mM NaCl, and fractions were collected. The protein-containing fractions determined by SDS-PAGE were pooled, and the protein concentration was measured at 280 nm (ε_280_ = 1490 M^−1^cm^−1^) using NanoDrop spectrophotometer (Thermo Fisher Scientific, Waltham, USA). Ammonium sulphate was added to the protein solution to obtain a final concentration of 0.3 mg/mL, and placed on a rocker for 30 min at 4 °C. Samples were then centrifuged at 20.000 x *g* for 35 min, at 4 °C, and precipitated protein was suspended in appropriate buffers based on the assembly reaction to be performed. An aliquot was injected into FPLC equipped with Superdex 75pg 60/300 or HiLoad 16/600 Superdex 75pg (Cytiva) columns.

### Tau (297-391) aggregations and seed productions

Tau seeds to be employed in seeded-aggregation experiments of Aβ42 were produced as described by Lovestam. S. *et al* ^24^. Assembly reactions were carried out in aliquots of 70 μL of purified tau (297–391) resuspended at 6 mg/mL. To produce PHF assembly, tau (297-391) was suspended in 10 mM PB, pH 7.2, containing 100 mM MgCl2, and 10 mM DTT, whereas CTE aggregates were obtained with tau297-391 suspended in 50 mM PB, pH 7.2, containing 100 mM NaCl, and 10 mM DTT. Unspecific fibrils were produced employing the tau suspended in the same solution used to obtain PHF assembly, added with heparin in a tau: heparin 4:1 ratio (w/w). Two μM ThT was added to each reaction mixture and the aggregation reactions were carried out at 37 °C in 96-well half-area low bind plates (Corning 3008), placed in a Tecan Infinite F200 equipped with 448 nm excitation and 485 nm emission filters. Samples were shaken using 200 r.p.m. the fluorescence was recorded every 5 min for 15 h. Fibrils obtained at this time point were subsequently bath sonicated for 5 min at room temperature and used to seed Aβ42 aggregation in cross-seeded aggregation experiments.

### ANP aggregation and seed production

Lyophilised synthetic ANP (GenScript, Piscataway, USA) was dissolved in 20 mM HEPES, pH 7.4, containing 1 mM NaCl, and centrifuged for 10 min at 20,800 x *g* at 4°C. The solution was diluted to 500 µM ANP in the same buffer and supplemented with 20 µM ThT^43^. Aggregations were performed at 37°C, applying a 600 r.p.m. linear shaking in 96-well half-area low bind plates (Corning 3008), placed in a Tecan Infinite F200 (Tecan) equipped with 448 nm excitation and 485nm emission filters. ANP fibrils obtained after 50 h were then bath sonicated for 5 min at room temperature to obtain seeds.

### Cross-seeded Aβ42 aggregations

Aβ42 cross-seeding aggregations with different tau seeds were performed, monitoring the aggregation of 4 µM monomeric Aβ42 in the absence or presence of 0 - 1 µM PHF, CTE, and disease-unrelated heparin seeds (expressed as fibril mass concentrations *M(0)_tau_*). Similar experiments were performed using 4 µM monomeric Aβ42 and 0.125-1µM ANP seeds. Cross-seeding aggregations were also performed using variable monomeric Aβ42 concentrations (3, 4, 5, 6, 7 µM) and 0.25 µM PHF and CTE tau seeds.

For all experimental settings, tau seeds produced as described above were diluted in the aggregation buffer (20 mM PB, pH 8, 200 µM EDTA, 0.02% NaN3, 5 mM NaOH, 2 µM ThT) and incubated for 1 h at room temperature before adding Aβ42 monomers. The aggregation reactions were followed for 15 h at 37°C in quiescent conditions. The fluorescence profiles were corrected for the seed contribution to the fluorescence signal, approximately linear in the timescale of the aggregation reaction. Each data set was analysed by fitting with the logistic function^37^:

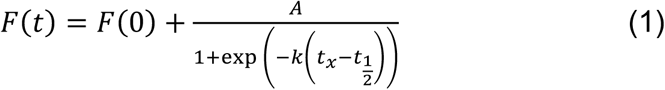

where *F(t)* and *F(0)* are the ThT fluorescence measured at each time point and at time zero. respectively, *A* is the amplitude of the fluorescence signal, *k* is the observed kinetic constant, and t_1/2_ is the half-time of the aggregation, corresponding to the time when 50% of monomers are found aggregated into amyloids^35^. t_lag_ was subsequently calculated as t_lag_ = t_1/2_ – 2/k ^37^. Microscopic kinetic analysis was performed using AmyloFit Platform (https://amylofit.com)^33^, global fitting to a model accounting for saturated multi-step secondary nucleation^39^ the kinetic curves of Aβ42 aggregations collected at increasing concentrations of Aβ42 (3 – 6 µM) incubated with 0.25 µM of PHF or CTE seeds. The elongation constant was assumed to be not affected by the presence of tau, to decrease the amount of degree of freedom for the fitting procedure. Saturation is described by an equilibrium constant (K_M_) correlated with the threshold concentration above which secondary nucleation saturates. Secondary nucleation in both self- and seeded conditions follows a second-order reaction (n_2_=2), and K_M_ is therefore m(0)^2^_threshold_. Control misfit was performed by global fitting to a secondary nucleation mechanism not accounting for saturation effects setting k_+_, k_2_ equal to the ones determined for Aβ42 self-aggregation. The kinetic parameters determine by global fitting were subsequently used to fit the kinetic traces obtained in cross-seeded reactions with 4µM Aβ42 and 0.125 – 1 µM PHF or CTE seed.

The primary nucleation constant k_n_ was group fitted for each seed concentration tested and used to calculate the λ values as: 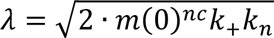

Enzyme-like behavior of tau seeds was determined adding 0.1 µM PHF and CTE seeds to the aggregation of 2 – 7 µM Aβ42, corresponding to 1:30 to 1:70 tau:Aβ42 molar ratios, optimized to determine the catalytic effect of tau fibrils. λ values were determined by global fitting as described before and fitted to a Hill kinetic equation (2) accounting for cooperativity mechanisms^31^.

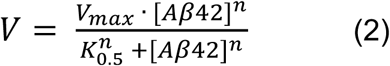

Where [Aβ42] is Aβ42 monomer concentration, V is the reaction velocity, V_max_ is the maximum velocity, K_0.5_ is the half-maximum concentration constant, and n is the Hill coefficient measuring the degree of cooperativity of substrate binding. If n =1 the equation is reduced to a Michaelis-Menten kinetic without cooperativity, n>1 positive cooperativity where the binding of one substrate to the catalytic sites facilitates the binding of other substrate molecules, whereas if n<1 is indicative of negative cooperativity.

### Immunoprecipitation experiments

Dynabeads M-280 Sheep anti-mouse IgG (Novex, Manchester, UK) were washed as described by the manufacturer’s instructions and coupled overnight, at 4°C, under mechanical agitation, with the murine monoclonal anti-Aβ, 1-16 Antibody (clone 6E10) (BioLegend, San Diego, USA). Functionalized beads were then added with the samples collected from Aβ self-aggregation, tau self-aggregation, and Aβ-tau cross-aggregation experiments, 0 and 15 h after the start of the aggregation reaction. After incubation for 2 h at 4°C under agitation, the tubes were placed on DynaMag (Novex, Manchester, UK), to separate the beads-containing pellet and the supernatant. The pellets were resuspended in 20 µL of loading buffer composed of 25 mM Tris solution containing 200 mM glycine, 0.2% SDS, 5% glycerol, and 0.025% bromophenol blue, and 10 µL was analyzed by immunoblotting using a 15% acrylamide gel and Western blotting. The membranes were blocked with 10 mM Tris-HCl solution, pH 7.5, containing 100 mM NaCl, 0.1% (v/v) Tween 20, 5% (w/v) low-fat dry milk powder, and 2% (w/v) bovine serum albumin, and incubated overnight with the anti-human Aβ42 rabbit polyclonal antibody (1:1000, Thermo Fisher Scientific), and the anti-human Tau rabbit polyclonal antibody (1:10,000, DAKO, Agilent Technologies Italia S.p.A., Cernusco sul Naviglio, Milano, Italy). Anti-rabbit IgG peroxidase conjugate (1:20,000, Sigma Aldrich) was used as a secondary antibody.

### TEM

Samples for negative-staining TEM continuous-carbon (2.5 µL) were directly applied to 300-mesh copper TEM grids (EMResolutions, Newcastle, UK), previously glow-discharged at 25 mA for one minute (GloQube Dual Chamber, Quorum Technologies, Lewes, UK), for 40 seconds to adhere. Grids were then negatively stained with 2.5 µL of 2 % (w/v) uranyl acetate for 40 seconds and air-dried for ≥ 2 minutes before either being stored or used immediately. Micrographs were acquired using a Talos F200X G2 TEM (Thermo Scientific, Waltham, USA) and a Ceta 4k × 4k CMOS camera. Manual measurements of cross-over distances and fibril widths were performed with Fiji platform^51^. Analysis of fibril crossover distance distributions were performed with one-sample and two-sample Kolmogorov– Smirnov (KS) tests^52^. One-sample Lilliefors-corrected KS tests were employed to evaluate if each dataset followed a normal distribution with normal distribution parameters estimated from the data. Two-sample KS tests were performed comparing empirical cumulative distribution functions (ECDFs) of different fibril populations and used to assess whether two distinct populations originated from the same underlying distribution. KS statistic D was defined as the maximum vertical difference between distributions, and significance was assessed at α=0.05. Sample size are: Aβ42 (n = 253), PHF-seeded Aβ42 (n = 239) and CTE-seeded Aβ42 (n =252). Analyses were performed using a custom Python script with the statsmodels module.

### Fibril Length Analysis

Micrographs of fibril populations used for analysis were taken at 36 000× magnification. Image stacks were demultiplexed and processed through a custom Python pipeline that integrates the SAM 2 (hiera-large) model (Meta)^53^, followed by additional filtering and morphological operations to generate individual binarized masks of fibril segments. Manual filtering was done to remove defective masks, including those which sampled background noise. For each mask, we assembled the covariance matrix of its foreground-pixel coordinates and identified the principal component (the major axis). Segment length was then determined by projecting all foreground pixels onto this axis and calculating the distance between their extreme projections. To determine segment width, we projected each foreground pixel onto the minor axis (orthogonal to the major axis), partitioned the major-axis projection into 1,000 evenly spaced windows, computed the peak-to-peak distance of the orthogonal projections within each window, and then reported the median of these slice widths. We converted pixel dimensions to nanometers using a calibration factor of 200 nm per 330.77 pixels, determined by running the analysis pipeline on a 200 nm scale bar.

### In silico docking analysis

In silico docking analysis was performed with HPEPDOCK^54^, starting from the PHF (PDB: 8Q8R) and CTE (PDB: 6NWP) fibril structures of tau and the Aβ42 sequence. 3D structures for the given peptide sequence were generated using the integrated MOPEP program. Default parameters were used to run the simulations, specifying the catalytic surface and surrounding residues, without introducing any energetic constraints. The region of tau spanning the residues 351-379 was chosen for docking and 10 fragments (**Supplementary Fig.4B)** of Aβ42 were employed for docking. One hundred poses for each fragment were generated and ranked based on the HPEPDOCK scoring function^55,56^. The highest-ranked prediction was employed for visualization and analysis with Chimera X software^57^.

### Experiments on SH-SY5Y neuroblastoma cells

SH-SY5Y human neuroblastoma cells (ATCC HTB-11, LGC Standards S.r.l., Sesto San Giovanni, Milano, Italy) were cultured at 37°C in Dulbecco’s Modification of Eagle’s Medium (DMEM) medium (Lonza, Basel, Switzerland) supplemented with 10% Fetal bovine serum (FBS, Gibco, Invitrogen, Waltham, MA, USA), 4 mM L-glutamine (Gibco), 50 U/mL penicillin and 50 μg/mL streptomycin, in humidified incubator with 5% CO_2_. Cytotoxic experiments were performed as described by Palmioli et al.^44^. Briefly, SH-SY5Y cells were seeded in 96-well plates (20,000 cells/200 μL/well) and incubated for 24 h at 37°C in a humidified atmosphere of 5% CO_2_. The medium was replaced with 200 μL/well of DMEM 1% FBS and cells were treated with 10 µM monomeric Aβ42 alone^44^ or together with 1.25 µM of PHF or CTE tau seeds suspended in the aggregation buffer. Control cells were treated with DMEM 1% FBS containing the aggregation buffer (Vehicle) or 1.25-5 µM of PHF or CTE tau seeds. Cell viability was evaluated 24 h later by adding 40 μL/well of 5 mg/mL 3-[4,5-dimethylthiazol-2-yl]-2,5-2,5-diphenyltetrazolium bromide (Sigma Aldrich). Two hours after incubation at 37°C, the medium was replaced with isopropanol acidified with 0.04 M, and the absorbance of the sample was spectrophotometrically determined at 570 nm, using the spectrophotometer Infinite M200 (Tecan, Männedorf, Switzerland).

### *C. elegans* experiments

Transgenic CL2120 *C. elegans*^46^, with genotype vIs14 [(pCL12) unc-54::beta 1-42 + (pCL26) mtl-2::GFP], expressing human Aβ42, its CL2122 control strain (dvIs15 [(pPD30.38) unc-54(vector) + (pCL26) mtl-2::GFP], and N2 Bristol UK wild-type worms were obtained from Caenorhabditis Genetics Center (CGC, University of Minnesota, Minneapolis, USA). Worms were grown at 20⁰C on solid Nematode Growth Medium (NGM) seeded with *E. coli* OP50 (CGC). Before the experiments, worms were synchronized by egg-laying and cultured for 72 h at 20°C on NGM plates spread with OP50 *E. coli* (CGC) as food. Worms were then collected by washing the plates with 1 mL M9 buffer (0.3% potassium dihydrogen phosphate, 0.6% sodium phosphate dibasic, 0.5% sodium chloride, and 1 M magnesium sulfate), CL2120 and CL2122 nematodes were administered with 20 and 100 µM PHF or CTE tau seed solutions (50 worms/50 μL). CL2120 control worms, CL2122, and N2 nematodes were administered 5 mM PBS, pH 7.4 (50 worms/50 μL). After 2 h of orbital shaking, worms were transferred onto fresh NGM plates seeded with fresh OP50 *E. coli*, put at 20°C, and their motility was determined 24 h and 48 h later by scoring the number of body bends per minute (body bends/min).

### Statistical analyses

No randomization was required for *C. elegans* experiments. All evaluations were done blind to the sample identity and treatment group. The data were analyzed using GraphPad Prism 10.2 software (San Diego, CA, USA) by Student’s *t*-test, one-way or two-way ANOVA, and Bonferroni’s or Tukey’s *post hoc* test. A *p*-value < 0.05 was considered significant.

