## Supplementary data for "Tau catalyzes amyloid-β aggregation in a fold-dependent manner"

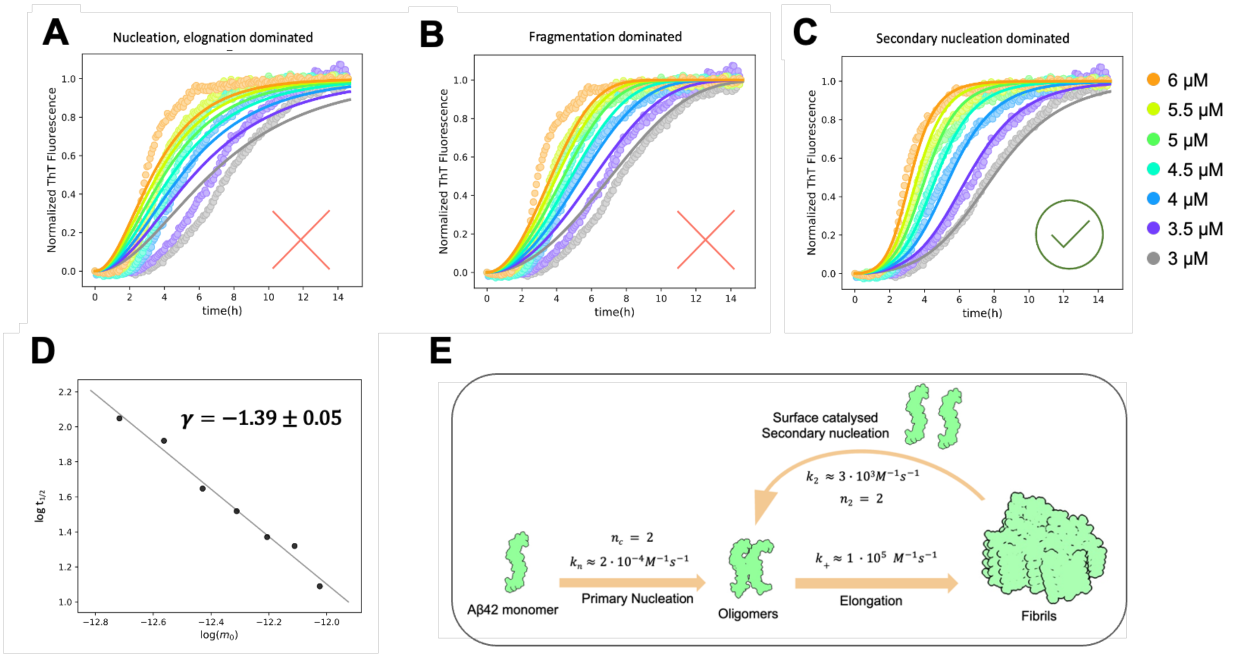


**Supplementary Figure 1. Microscopic aggregation mechanism of Aβ42. A-C.** Representative normalized aggregation curves obtained by monitoring ThT fluorescence for different Aβ42 monomer concentrations ($m_0$) at 37°C in quiescent conditions in 20 mM PB, pH 8, 200 μM EDTA, 0.02% $NaN_3$, 5 mM NaOH, 2 μM ThT. **A.** Misfit with a nucleation + elongation kinetic model, not accounting for secondary pathways. **B.** Misfit with a fragmentation-dominated model. **C.** The best fit resulting from the fitting with a secondary nucleation model. **D.** Estimation of the scaling exponent by plotting $t_{1/2}$ *vs.* $m_0$ in a double logarithmic plot confirms the predominance of secondary nucleation in guiding the aggregation of Aβ42. **E.** Schematic showing the microscopic parameters underlying the overall Aβ42 aggregation. Monomers interact to form critical elongation-prone nuclei through primary nucleation, and fibrils are subsequently produced by elongation. Surface-catalyzed secondary nucleation is determined to be responsible for the exponential production of amyloid fibrils.

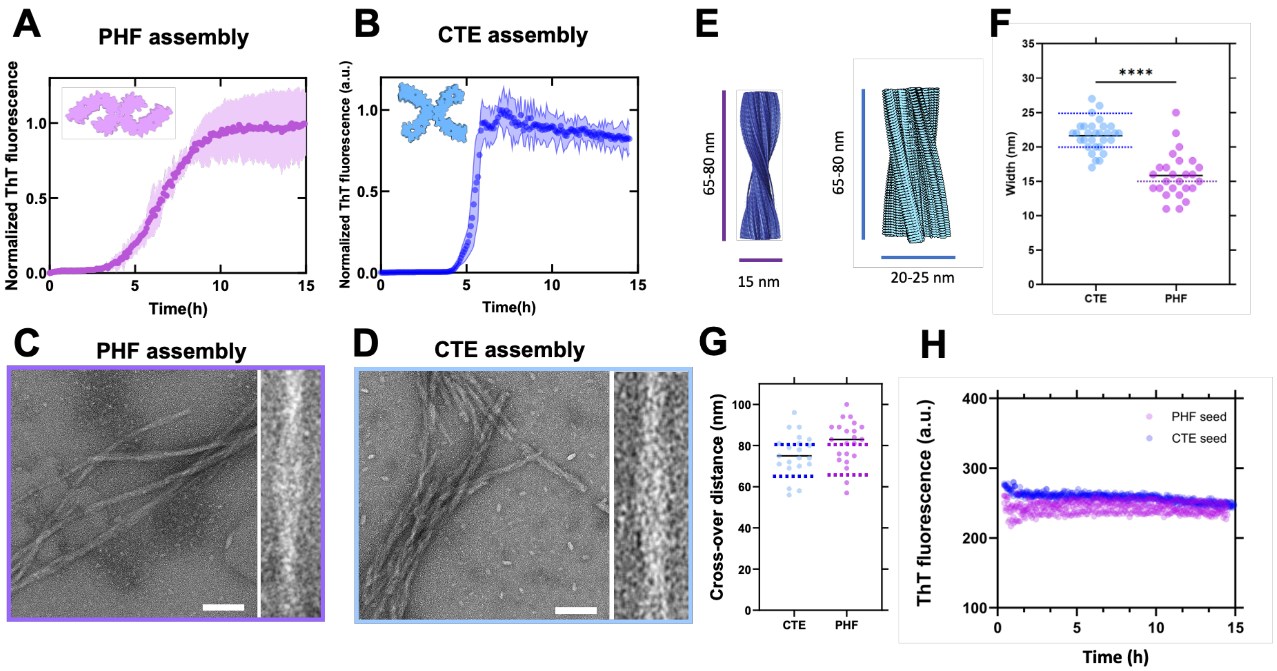


**Supplementary Figure 2. Tau297-391 aggregation in PHF and CTE assembly conditions.** Representative ThT fluorescence profiles of tau297-391 aggregation in **A.** PHF assembly conditions (10 mM phosphate buffer at pH 7.2, 100 mM $MgCl_2$, and 10 mM DTT) and **B**. CTE assembly conditions (50 mM phosphate buffer at pH 7.2, 100 mM NaCl, and 10 mM DTT). Reactions were carried out for 15 h using 200 r.p.m. orbital shaking at 37°C, monitoring the fluorescence of 2 µM ThT. The first 12-15 hours are shown. Each point is the mean ± SEM of at least two replicates. **C, D.** Representative negative staining TEM images of fibrils produced respectively in PHF and CTE assembly conditions. Detailed views of fibril morphologies are on the right of each micrograph. Scale bars are 100 nm. **E.** Schematic showing characteristic width and cross-over distance of PHF (blue) (from Fitzpatrick, A., et al. *Nature* 547, 185–190 (2017)) and CTE (green) (PDB: 6NWP) fibrils. **F.** Measured fibril widths of tau fibrils produced in CTE and PHF assembly reactions compared to values reported in literature for each type (dashed lines). Black lines are experimental means (N > 30). ****$p < 0.0001$, Student's t-test. **G.** Measured fibril cross-over distances of tau fibrils produced in CTE and PHF assembly reactions compared to values reported in literature for each type (dashed lines). Black lines are experimental means (N > 30). **H.** Representative ThT fluorescence profiles of 0.25 µM tau297-391 seeds with PHF (purple) and CTE (blue) in cross-seeded aggregation conditions without Aβ42. The seeds displayed a linear behaviour in the time scale of cross-seeded aggregations (15 h). In the present study, we treated seed fluorescence as baseline and they were subtracted for all Aβ42 cross-seeded aggregations.

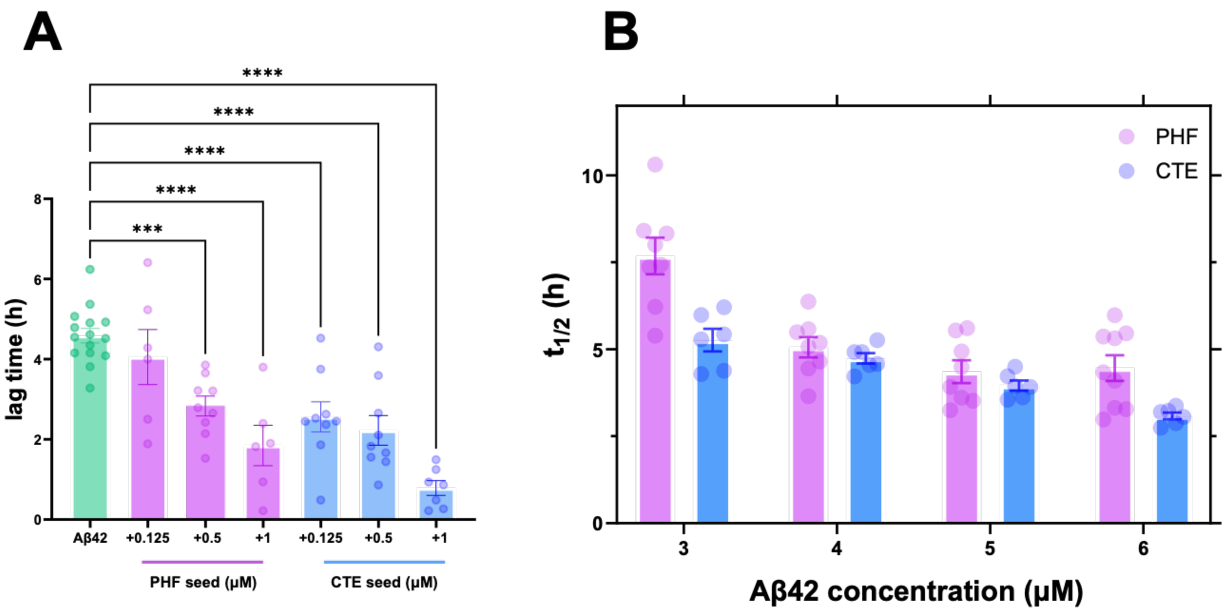


**Supplementary Figure 3. Effect of tau seeds on Aβ42 aggregation. A.** Increasing concentrations of tau seeds accelerate Aβ42 aggregation, reducing the lag time of the reaction in a dose-dependent manner. The green bar is the lag time of 4μM Aβ42 self-aggregation. The lag time of seeded aggregation with 0.125, 0.5, and 1 μM PHF (purple) and CTE (blue) is also reported. Data are mean ± SEM of experimental data (reported as dots) from at least three independent experiments. Statistical differences between seeded and unseeded aggregation were assessed with one-way ANOVA and Bonferroni's *post hoc* test analysis (***$p<0.001$ and ****$p<0.0001$). **B.** Half-time of seeded aggregations. Effect of 0.25 μM PHF (purple) and CTE (blue) on 3-6 μM Aβ42. Data are mean ± SEM (N=3) of experimental data (reported as dots) from at least three independent experiments.

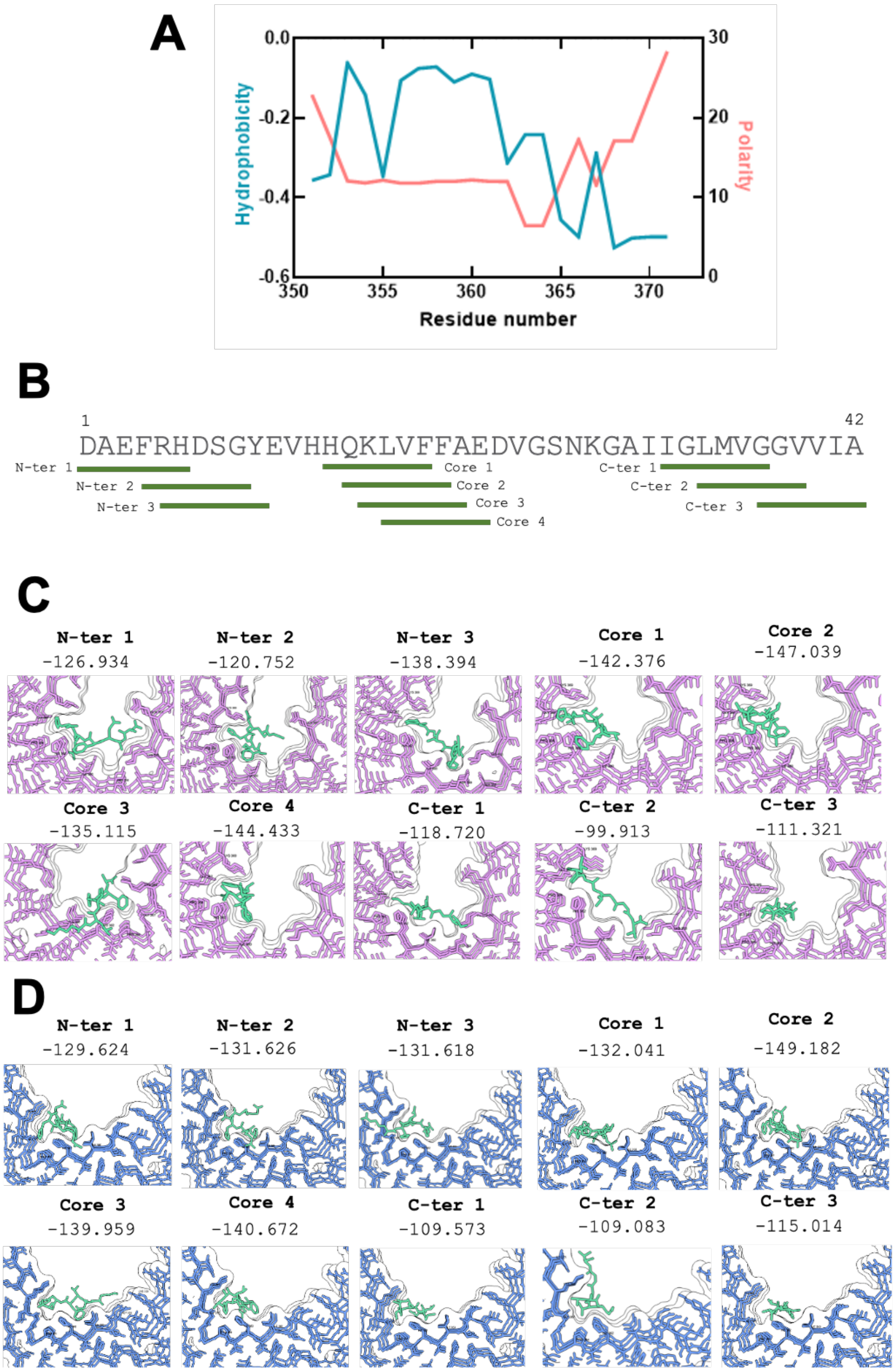


**Supplementary Figure 4. Docking analysis of Aβ42 segments on tau fibrils. A.** Hydrophobicity (Sweet R.M., Eisenberg D. J. Mol. Biol. 171:479-488(1983) and polarity (Zimmerman JM, Eliezer N, Simha R. J. Theor. Biol. 21:170-201(1968) profiles of the tau surface employed in docking analysis. The N-terminal portion presents high hydrophobicity, while the C-terminal region is highly polar, constituting a suitable constrained environment for the heterotypic nucleation of Aβ42. In CTE fibrils, this surface is more exposed to the solvent, consistent with an increased reactivity. **B.** Aβ42 primary sequence, in green are indicated the hexapeptides employed for docking**.** Docking analysis on **C.** PHF and **D.** CTE tau fibrils. Highest-scoring predictions are shown for each hexapeptide. Values are scores assigned by HPEPDOCK to each prediction for ranking.

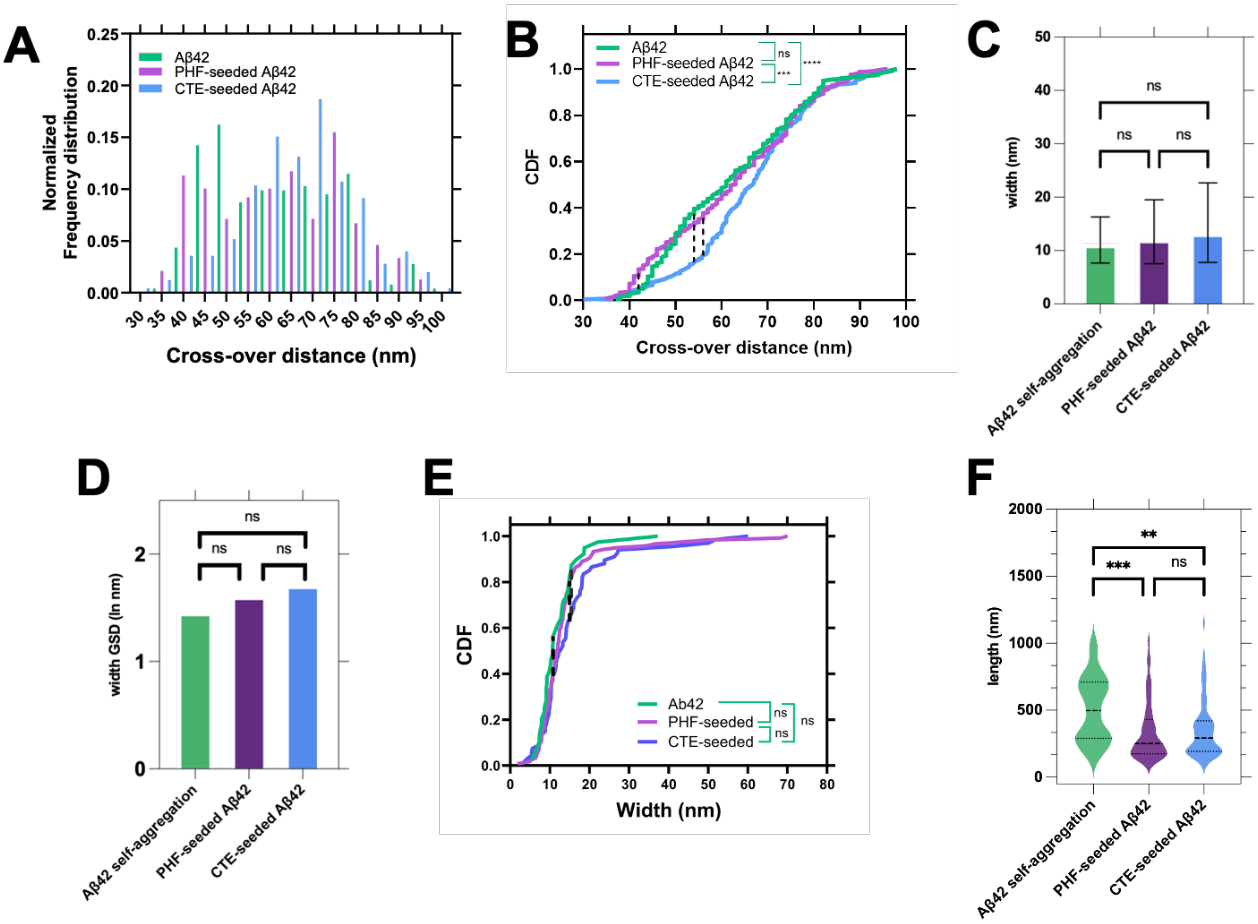


**Supplementary Figure 5. Morphological characterization of Aβ42 fibrils. A.** Frequency distribution of crossover distances of Aβ42 fibrils. CTE-seeded Aβ42 fibrils are distributed around an average cross-over distance of 66 nm, whereas other fibrils present random distribution in the 30-100 nm range. **B.** CDFs of cross-over distances for PHF- seeded (purple), CTE-seeded (blue), and self-aggregated (green) Aβ42 fibrils. Pairwise two-sample Kolmogorov–Smirnov (KS) tests were performed: Aβ42 *vs.* PHF-seeded Aβ42 ($D_{0.05}$ = 0.122, D = 0.086, not significant), Aβ42 *vs.* CTE-seeded Aβ42 ($D_{0.05}$ = 0.121, D = 0.229, $p < 0.0001$), and PHF-seeded *vs.* CTE-seeded Aβ42 ($D_{0.05}$ = 0.123, D =0.182, $p < 0.001$). D represents the maximum vertical separation between CDFs (dashed lines), $D_{0.05}$ is the critical value at significance 0.05. **C.** Measurements of fibril width of Aβ42 fibrils formed in self-aggregation, PHF- and CTE-seeded conditions conducted using a custom Python pipeline that integrates the SAM 2 (hiera-large) model. Data are mean ± SD (N > 100). Statistical analysis was performed with one-way ANOVA and Bonferroni's *post hoc* test (ns = not significant). **D.** Geometric standard deviation (GSD) of width measurements analysed with the Levene test, confirming no significant differences in variance between different populations. **E.** CDFs of fibril width for PHF- seeded (purple), CTE-seeded (blue), and self-aggregated (green) Aβ42 fibrils. Pairwise two-sample Kolmogorov–Smirnov tests were performed: Aβ42 *vs*. PHF-seeded Aβ42 ($D_{0.05}$ = 0.251, D = 0.174, not significant), Aβ42 *vs.* CTE-seeded Aβ42 ($D_{0.05}$ = 0.274, D = 0.215, not significant), and PHF-seeded *vs.* CTE-seeded Aβ42 ($D_{0.05}$ = 0.208, D = 0.178, not significant). D represents the maximum vertical separation between CDFs (dashed lines), $D_{0.05}$ is the critical value at significance 0.05. **F.** Fibril lengths analysis of Aβ42 amyloids produced in the three different aggregation conditions. Statistical differences between seeded and unseeded aggregation were assessed with one-way ANOVA and Bonferroni's *post-hoc* test (**$p$ <0.01; ***$p$<0.001).

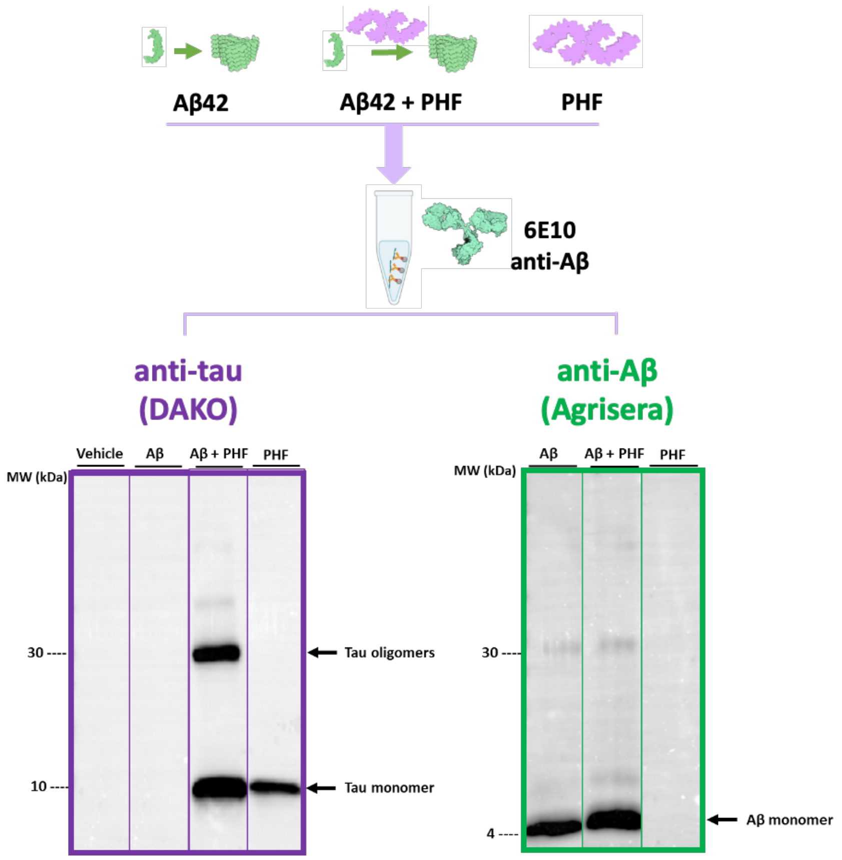


**Supplementary Figure 6. Tau-Aβ42 interaction demonstrated by immunoprecipitation.** Aβ42 fibrils (4 μM) produced 15 h after aggregation in 0.5 μM PHF-seeded conditions were immunoprecipitated with anti-Aβ mouse monoclonal 6E10 antibody. Unseeded Aβ42 fibrils (4 μM) and PHF seeds (0.5 μM) were used as positive controls. The aggregation buffer was employed as a negative control. Aβ42 and tau in the post-immunoprecipitation pellet, suspended in loading buffer, were analyzed by 15% SDS-PAGE. Ten μL were loaded in each gel lane and immunoblotted with an anti-tau rabbit antibody (DAKO) or an anti-human Aβ42 rabbit polyclonal antibody (Agrisera). Anti-rabbit IgG peroxidase conjugate was used as a secondary antibody.

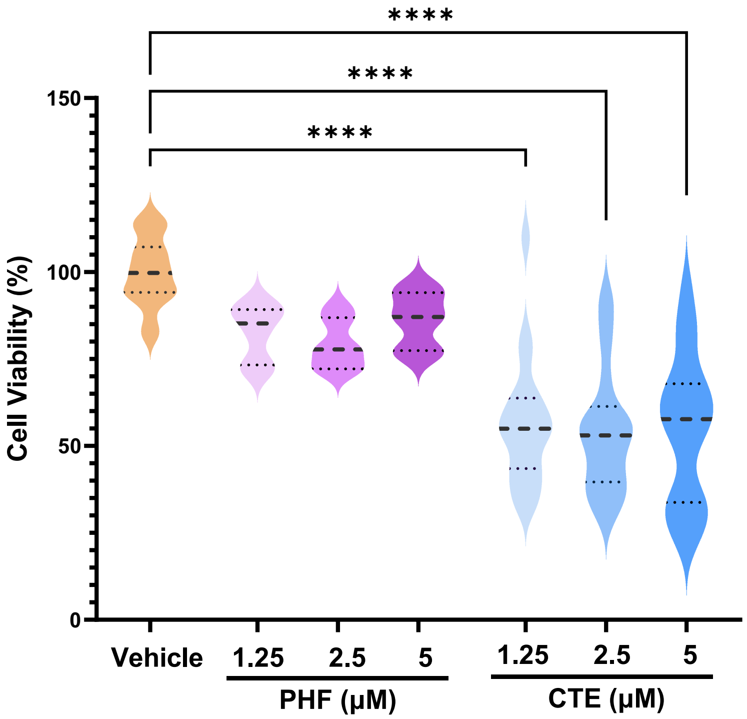


**Supplementary Figure 7. Effect of different tau seed concentrations on the viability of SH-SY5Y cells.** SHSY-5Y neuroblastoma cells were treated with 1.25-5 µM PHF or CTE seeds, and the cell viability was determined after 24 h. Control cells were treated with the medium alone, containing the aggregation buffer (Vehicle). Data are expressed as the mean ± SD of % viability compared to Vehicle-treated cells (N = 10). ****$p$<0.0001 according to one-way ANOVA and Bonferroni *post hoc* test.

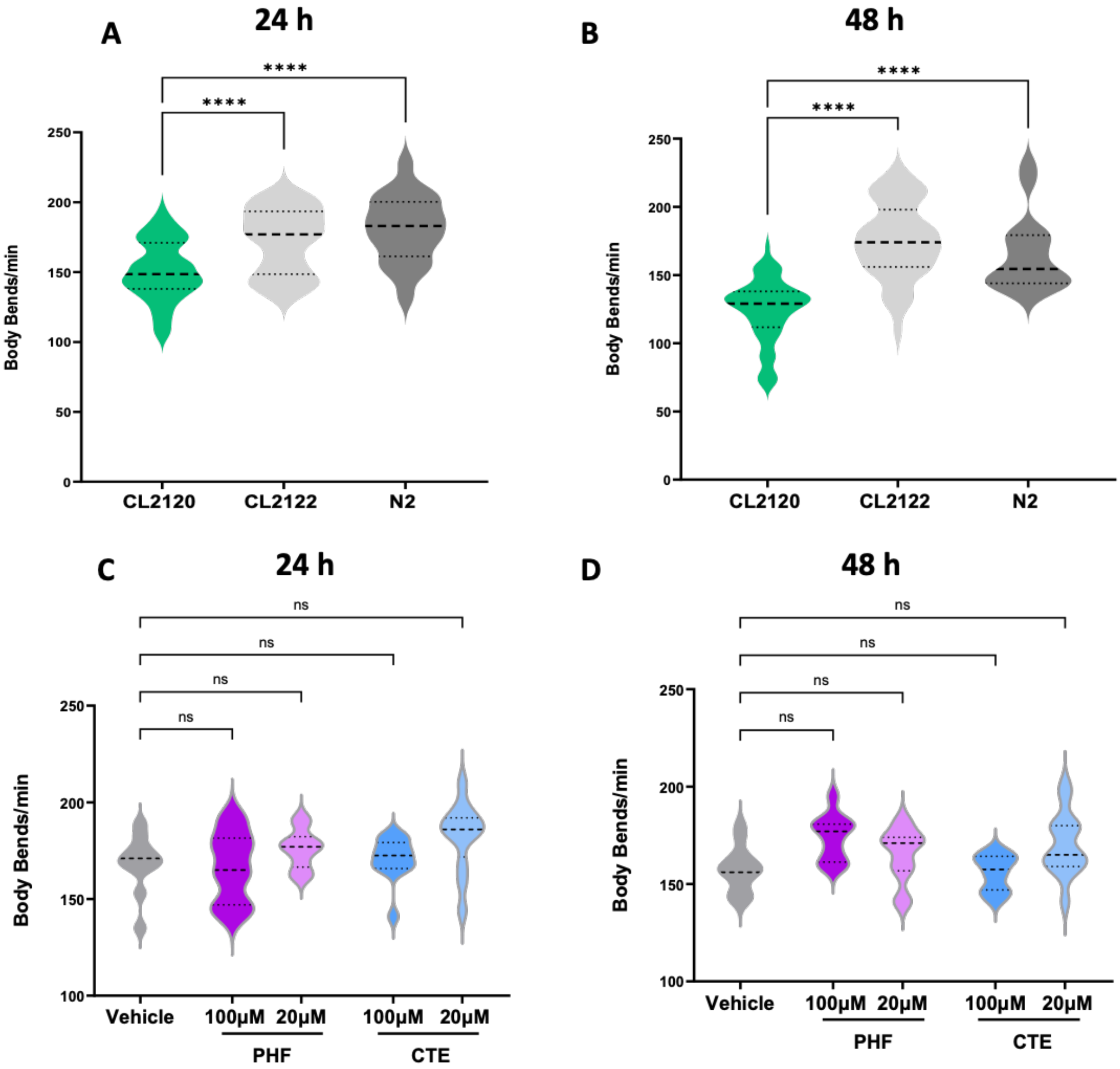


**Supplementary Figure 8. Effect of Aβ42 expression on the motility of the CL2120 transgenic strain and impact of tau polymorphs on CL2122.** The motility of synchronized transgenic CL2120 worms expressing Aβ42, and of CL2122 and N2 control worms, was determined at the **A.** L4 larval stage and the **B.** first day of adulthood, corresponding to the time points of 24 h and 48 h. The motility of synchronized transgenic CL2122 control worms was determined at the **C.** L4 larval stage and the **D.** first day of adulthood, corresponding to 24 h and 48 h after tau seeds administration. Data are the mean ± SEM (N = 30) from 3 independent experiments. *****p*<0.0001, ns = not significant according to one-way ANOVA and Bonferroni *post hoc* test.

**A**

| PHF-seeded Aβ42 conc. variable | Multi-step Secondary Nucleation | Secondary Nucelation |
|---|---|---|
| $k_n$ $(M^{-1}h^{-1})$ | 3.5 | 1.9 |
| $n_c$ | 2 | 2 |
| $k_+$ $(M^{-1}h^{-1})$ | $3.5 \cdot 10^8$ | $3.5 \cdot 10^8$ |
| $k_2$ $(M^{-2}h^{-1})$ | $4.6 \cdot 10^8$ | $1 \cdot 10^7$ |
| $n_2$ | 2 | 2 |
| $K_M$ $(M^{n_2})$ | $6.1 \cdot 10^{-14}$ | - |
| MRE | 0.0014 | 0.0057 |

| CTE-seeded Aβ42 conc. variable | Multi-step Secondary Nucleation | Secondary Nucelation |
|---|---|---|
| $k_n$ $(M^{-1}h^{-1})$ | 3.7 | 6.8 |
| $n_c$ | 2 | 2 |
| $k_+$ $(M^{-1}h^{-1})$ | $3.5 \cdot 10^8$ | $3.5 \cdot 10^8$ |
| $k_2$ $(M^{-2}h^{-1})$ | $1 \cdot 10^{11}$ | $1 \cdot 10^7$ |
| $n_2$ | 2 | 2 |
| $K_M$ $(M^{n_2})$ | $1.3 \cdot 10^{-15}$ | - |
| MRE | 0.0031 | 0.0089 |

**B**

| | $k_n$ $(M^{-1}h^{-1})$ | $n_c$ | $k_+$ $(M^{-1}h^{-1})$ | $k_2$ $(M^{-2}h^{-1})$ | $n_2$ | $K_M$ $(M^{n_2})$ | MRE |
|---|---|---|---|---|---|---|---|
| Aβ42 | 0.78 | 2 | $3.5 \cdot 10^8$ | $4.6 \cdot 10^8$ | 2 | $2.02 \cdot 10^{-13}$ | 0.0016 |
| + 0.125μM PHF | 1.32 | 2 | $3.5 \cdot 10^8$ | $4.6 \cdot 10^8$ | 2 | $2.02 \cdot 10^{-13}$ | 0.0016 |
| + 0.25μM PHF | 1.97 | 2 | $3.5 \cdot 10^8$ | $4.6 \cdot 10^8$ | 2 | $2.02 \cdot 10^{-13}$ | 0.0016 |
| + 0.5μM PHF | 5.58 | 2 | $3.5 \cdot 10^8$ | $4.6 \cdot 10^8$ | 2 | $2.02 \cdot 10^{-13}$ | 0.0016 |
| + 1 μM PHF | 6.96 | 2 | $3.5 \cdot 10^8$ | $4.6 \cdot 10^8$ | 2 | $2.02 \cdot 10^{-13}$ | 0.0016 |
| Aβ42 | 0.60 | 2 | $3.5 \cdot 10^8$ | $1 \cdot 10^{11}$ | 2 | $1.3 \cdot 10^{-15}$ | 0.0013 |
| + 0.125μM CTE | 2.00 | 2 | $3.5 \cdot 10^8$ | $1 \cdot 10^{11}$ | 2 | $1.3 \cdot 10^{-15}$ | 0.0013 |
| + 0.25μM CTE | 5.17 | 2 | $3.5 \cdot 10^8$ | $1 \cdot 10^{11}$ | 2 | $1.3 \cdot 10^{-15}$ | 0.0013 |
| + 0.5μM CTE | 7.57 | 2 | $3.5 \cdot 10^8$ | $1 \cdot 10^{11}$ | 2 | $1.3 \cdot 10^{-15}$ | 0.0013 |
| + 1 μM CTE | 16.69 | 2 | $3.5 \cdot 10^8$ | $1 \cdot 10^{11}$ | 2 | $1.3 \cdot 10^{-15}$ | 0.0013 |

**Supplementary Table 1. A.** Global fitting analysis of kinetic curves collected at increasing Aβ42 concentrations (3- 6 μM) in the presence of 0.25 μM PHF (upper table) and CTE (bottom table). Normalized ThT curves were fitted to a multi-step secondary nucleation model, resulting in the best fit. $k_n$, $k_2$ and $K_M$ were globally fitted, $n_c$ and $k_+$ were set as global constants as determined for Aβ42 self-aggregation. $n_2$ was initially globally fitted, resulting in values around 2; it was then kept as a global constant to minimize the variables in the global fitting procedure. Global fit with a secondary nucleation model, neglecting the multi-step behaviour, imposing $k_+$, $k_2$, $n_c$, $n_2$ as global constants as for self-aggregation, resulted in a misfit for both CTE- and PHF-seeded cases. **B.** Global fitting analysis of kinetic curves collected at increasing concentrations of tau seeds. Kinetic parameters extrapolated in the previous fitting were used as global constants, resulting in a good fit. $k_n$ was group fitted for each seed concentration to determine the initial rate of fibril formation as described in the main text. In the PHF-seeded case, $K_M$ needed to be set as a variable; we provided the value obtained in the previous fitting as a starting guess. The value obtained still aligns with a saturation effect occurring at $\sqrt{K_M} < 3\mu M$.
